# Intratumoral Resident Microbes Directly Participate in Resistance to EGFR-TKI Targeted Drug Therapy through Metabolism in NSCLC

**DOI:** 10.1101/2025.03.12.642829

**Authors:** Huan Wang, Jiarui Zhang, Jiaxun Zhang, Wang Hou, Jing Zhou, Maolin Li, Xuyan Liu, Yangqian Li, Fei Fu, Panwen Tian, Weiming Li, Dan Liu, Dan Xie

**Author notes:** These authors contributed equally (Huan Wang, Jiarui Zhang, Jiaxun Zhang, Wang Hou).

## Abstract

EGFR-TKIs targeted therapy represents a cornerstone in the treatment of non-small cell lung cancer (NSCLC). Nonetheless, a subset of patients inevitably develops acquired resistance to EGFR-TKIs, posing a significant clinical challenge. An increasing number of evidence has documented the presence of microbial communities within tumor microenvironments, but the potential interplay between tumor-resident microbiota and resistance to EGFR-TKIs remains poorly understood. Here, we report that prolonged EGFR-TKIs treatment markedly alters the diversity of tumor-resident microbiota, characterized by a notable enrichment of *Bacillus mycoides* and *Pseudomonas veronii*. In vitro experiments revealed that co-culture of gefitinib with either *B. mycoides* or *P. veronii* significantly attenuated its tumor-suppressive efficacy, both in vitro and in vivo. Mechanistically, gefitinib underwent extensive biotransformation in the presence of these bacteria, yielding redox derivatives and conjugation products, which collectively diminished its antitumor activity. Specifically, *B. mycoides* metabolized gefitinib into B.mPM9-C33 H45 Cl F N3 O4, while *P. veronii* generated P.vPM2-C24 H29 Cl N4 O3, both of which were consistently detected in in vivo and in vitro models. Furthermore, heterologous expression of maltose acetyltransferase from *B. mycoides* in *E. coli BL21* conferred the capacity to metabolize gefitinib, underscoring the enzymatic basis of this metabolic conversion. Collectively, these findings demonstrate that *B. mycoides* and *P. veronii* drive tumor resistance to EGFR-TKIs through metabolic inactivation of the drug, unveiling a previously unrecognized role of tumor-resident microbiota in modulating therapeutic responses and providing novel insights into the mechanisms underlying EGFR-TKIs resistance in NSCLC.

## Introduction

The human microbiota has emerged as nonnegligible components of the tumor microenvironment, providing novel perspectives on cancer biology. While earlier research predominantly focused on oral and gut microbiota due to their easier sample accessibility, direct disease associations and high microbial density [1–3], recent advances have revealed microbial presence even in traditionally “sterile” organs. Mounting evidence demonstrates that tumor-resident bacteria persist across diverse malignancies including breast, lung, ovarian, pancreatic, melanoma, bone, and brain cancers [4], fundamentally expanding our understanding of host-microbe interactions in cancer. These intratumoral microbes exhibit remarkable functional versatility. Some residential bacteria invade tumor cells and travel through the circulation system along with the cancer cells and play critical roles in metastatic colonization by modulating the host-cell actin network and promoting cell survival against fluid shear stress in the circulation of breast cancer [5], and Lactic acid bacteria within the tumor microenvironment can alter tumor metabolism and lactate signaling pathways, causing chemotherapy and radiation resistance in cervical cancer [6]. In pancreatic cancer, bacterial species harboring long-form cytidine deaminase within tumors have been causing chemotherapy resistance [7]. This paradigm shift underscores the need to investigate how tumor-colonizing microbes may directly shape therapeutic outcomes.

In parallel, the management of EGFR-mutant non-small cell lung cancer (NSCLC) has been transformed by EGFR tyrosine kinase inhibitors (TKIs), which achieve superior progression-free survival compared to chemotherapy [8–13]. While third-generation inhibitors like Osimertinib have revolutionized care for patients with T790M mutation and brain metastases[14–16], acquired resistance inevitably emerges within 9-12 months of treatment initiation due to either EGFR-dependent mechanisms (e.g., C797S mutations) or alternative bypass pathways signaling activation or both [17–19]. Despite extensive characterization of tumor cell-autonomous resistance mechanisms, critical gaps remain in understanding microenvironmental contributors to TKI resistance.

Notably, while pharmacological interventions are known to reshape gut microbial communities [20–22], the potential crosstalk between long-term TKI exposure and tumor-resident microbiota remains unexplored. This intersection presents a compelling scientific opportunity: Given that pulmonary tumors harbor distinct bacterial communities and that microbial metabolites can influence drug efficacy across cancer types [23, 24]. we hypothesize that tumor-resident bacteria play a potential role in EGFR TKI resistance. Elucidating these interactions could unveil novel mechanisms of therapeutic failure and microbiota-targeted strategies to overcome resistance in NSCLC. In this study, we aimed to investigate the potential link between intratumoral microbiota and resistance to EGFR-TKIs in NSCLC. We collected tissue samples from 128 patients with NSCLC, encompassing those prior to targeted EGFR treatment and those developed resistance after prolonged EGFR-TKIs therapy. Through 16S rRNA long-read third-generation sequencing, we discovered significant variations in microbial α-diversity and β-diversity within the same patient before and after resistance emergence, with a notable reduction in microbial diversity in the tissue post-resistance. The comparative analysis of unpaired samples revealed two significantly enriched bacterial species in resistant tumors: *B. mycoides* and *P. veronii*. Through in vitro experiments and mouse models, we demonstrated that the presence of *B. mycoides* and *P. veronii* can assist NSCLC in tolerating Gefitinib. Mechanistically, we demonstrated that both in vivo and in vitro, *B. mycoides* can metabolize gefitinib to B.mPM9-C33 H45 Cl F N3 O4, while *P. veronii* can metabolize gefitinib to P.vPM2-C24 H29 Cl N4 O3 with the oxidoreductase and maltose acetyltransferase identified as the key enzymes involved in metabolizing gefitinib which cause acquired resistance to this drug. In summary, our study uncovered a previously unrecognized role of NSCLC tissue-resident microbiota in promoting resistance to EGFR-TKI targeted drug therapy and highlights the potential for microbiota targeted intervention to overcome therapeutic resistance.

## Results

### Long-term targeted drug therapy has changed the abundance of resident bacteria in lung cancer

The emerging evidence of tumor-resident microbiota in human malignancies prompted us to investigate their potential involvement in EGFR-TKI resistance mechanisms. In this study, we enrolled a cohort of 144 patients with histologically confirmed, locally advanced NSCLC (Table S1). Among these, pretreatment tumor specimens were obtained from 52 treatment-naïve patients prior to EGFR-TKI administration, while post-treatment biopsies were collected from 76 patients following the development of acquired resistance. Notably, we secured matched pre- and post-resistance tumor samples from 7 patients, providing a unique opportunity for longitudinal analysis (Fig. 1a; Table S1). To characterize the intratumoral microbial landscape, we performed full-length 16S rRNA gene sequencing using the PacBio platform, analyzing tumor tissues from seven NSCLC patients who underwent EGFR-TKI therapy, with matched pre-treatment and post-resistance specimens (n=14) (Table S1). Comparative microbiome analysis revealed significant alterations in both α-diversity (Shannon index) and β-diversity between pretreatment and post-resistance samples (Fig. 1b, c). Through linear discriminant analysis (LDA) effect size (LEfSe) measurements, we identified five bacterial species showing significant enrichment in post-resistance tissues: *Pseudomonas reactans*, *P.veronii*, *Lactobacillus iners*, *B. mycoides*, and *Neisseria sicca* (Fig. 1d, e; Extended Data Fig. 1a, b). Quantitative assessment of these tumor-associated microbiota in an expanded cohort of unmatched samples (Table S1) demonstrated that *B. mycoides* emerged as the predominant bacterial species in EGFR-TKI-resistant tumors (Extended Data Fig. 1c, d). Furthermore, both *B. mycoides* and *P. veronii* showed consistent enrichment patterns in post-resistance tissues across the unmatched sample set (Fig. 1e). Based on these findings, we focused our subsequent mechanistic investigations on the potential roles of *B. mycoides* and *P. veronii* in mediating EGFR-TKI resistance.

**Fig 1.**
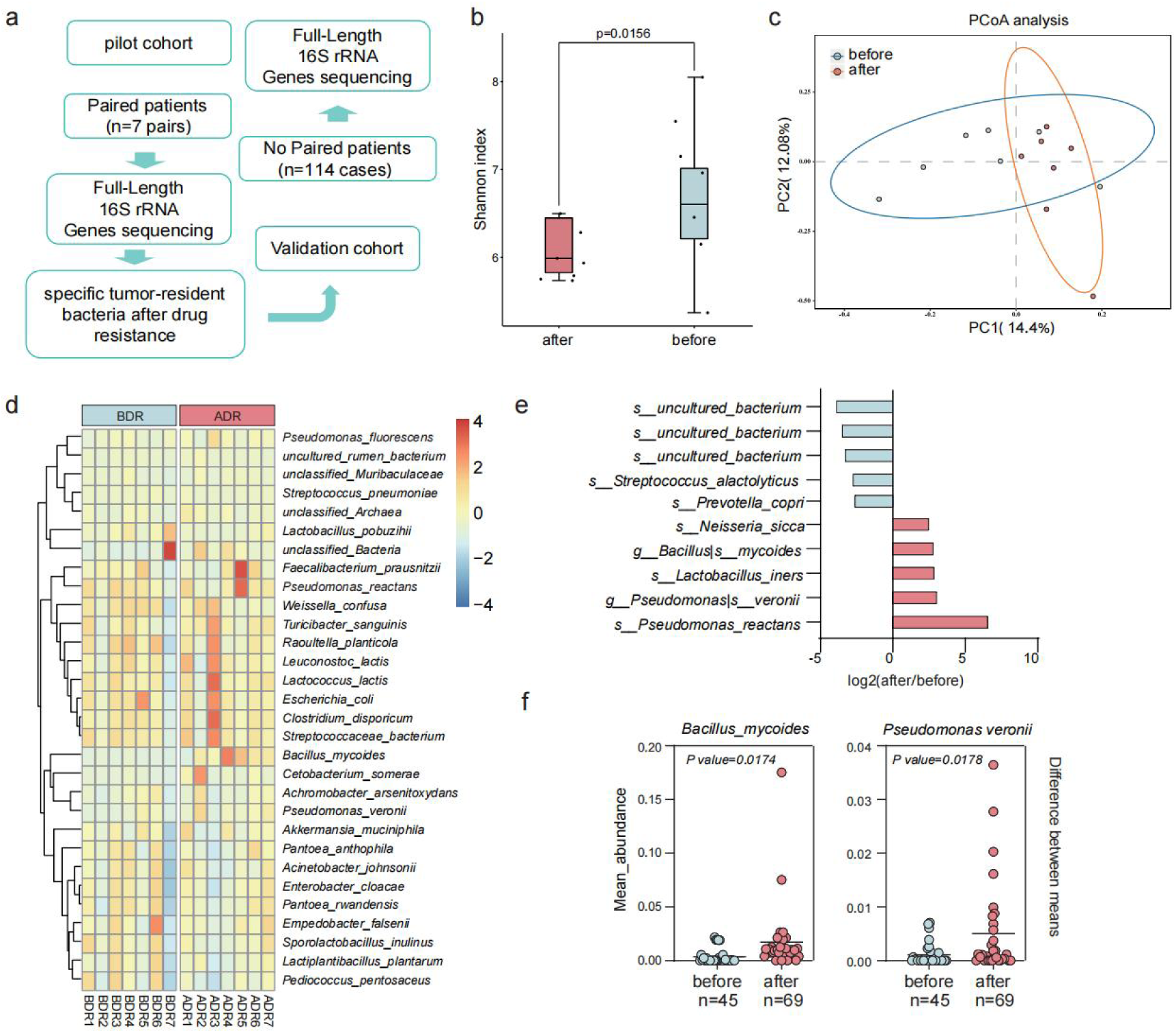
EGFR-TKIs therapy impacts the abundance of resident bacteria and specific bacterial taxa (*B. mycoides* and *P. veronii*) in lung cancer. a, Study design and clinical cohort characteristics; pilot cohort, paired samples of the same patient before TKIS treatment and after treatment resistance (n=7). Validation cohort, unpaired 114 samples, among which 45 samples are before drug resistance and 69 samples are after drug resistance. **b,c**,The microbial abundances in all samples were detected by full-length 16S rRNA gene sequencing technique of the PacBio platform. The alpha diversity (**b**) and PCoA analysis (**c**) of tumor-resident bacteria in pilot cohort. Mann ‒ Whitney test (**b**). **d,** Species abundance heatmap reveals differential tumor-resident microbial profiles between pre- and post-drug resistance in paired patient samples. **e,** Comparative bar plot identifies significantly altered bacterial species in pilot cohort following acquisition of drug resistance. **f,** Two bacterial taxa demonstrate significant enrichment associated with acquired drug resistance in the validation cohort. Among the 69 samples analyzed for post-drug resistance, 53 exhibited resistance to first-generation KTI drugs, with bacterial detection frequencies of 39.6% (21/53) for *B. mycoides* and 64.2% (34/53) for *P. veronii.* 13 samples demonstrated resistance to third-generation KTI drugs, with lower bacterial detection frequencies of 15.4% (2/13) for *B. mycoides* and 38.5% (5/13) for *P. veronii*.

### *B. mycoides* and *P. veronii* assist tumor cells to tolerate Gefitinib

To elucidate the potential causal relationship between *B. mycoides, P. veronii,* and EGFR-TKI resistance, we conducted a series of in vitro experiments to assess the impact of these bacterial species on gefitinib efficacy. Bacterial suspensions of *B. mycoides* and *P. veronii* were individually co-incubated with gefitinib for 2 hours, followed by filtration to remove bacterial cells, and the resulting solutions were subsequently co-cultured with either gefitinib-sensitive PC9 or gefitinib-resistant A549 cell lines. In both cellular models, pre-treatment of gefitinib with *B. mycoides* or *P. veronii* significantly attenuated its antiproliferative activity. Quantitative analysis revealed that gefitinib pre-exposed to *B. mycoides* (Fig. 2a,b and Extended Data Fig. 2a,c) or *P. veronii* (Fig. 2c,d and Extended Data Fig. 2b,d) exhibited diminished capacity to induce cell death compared to untreated gefitinib controls, as evidenced by increased cellular proliferation rates.

**Fig 2.**
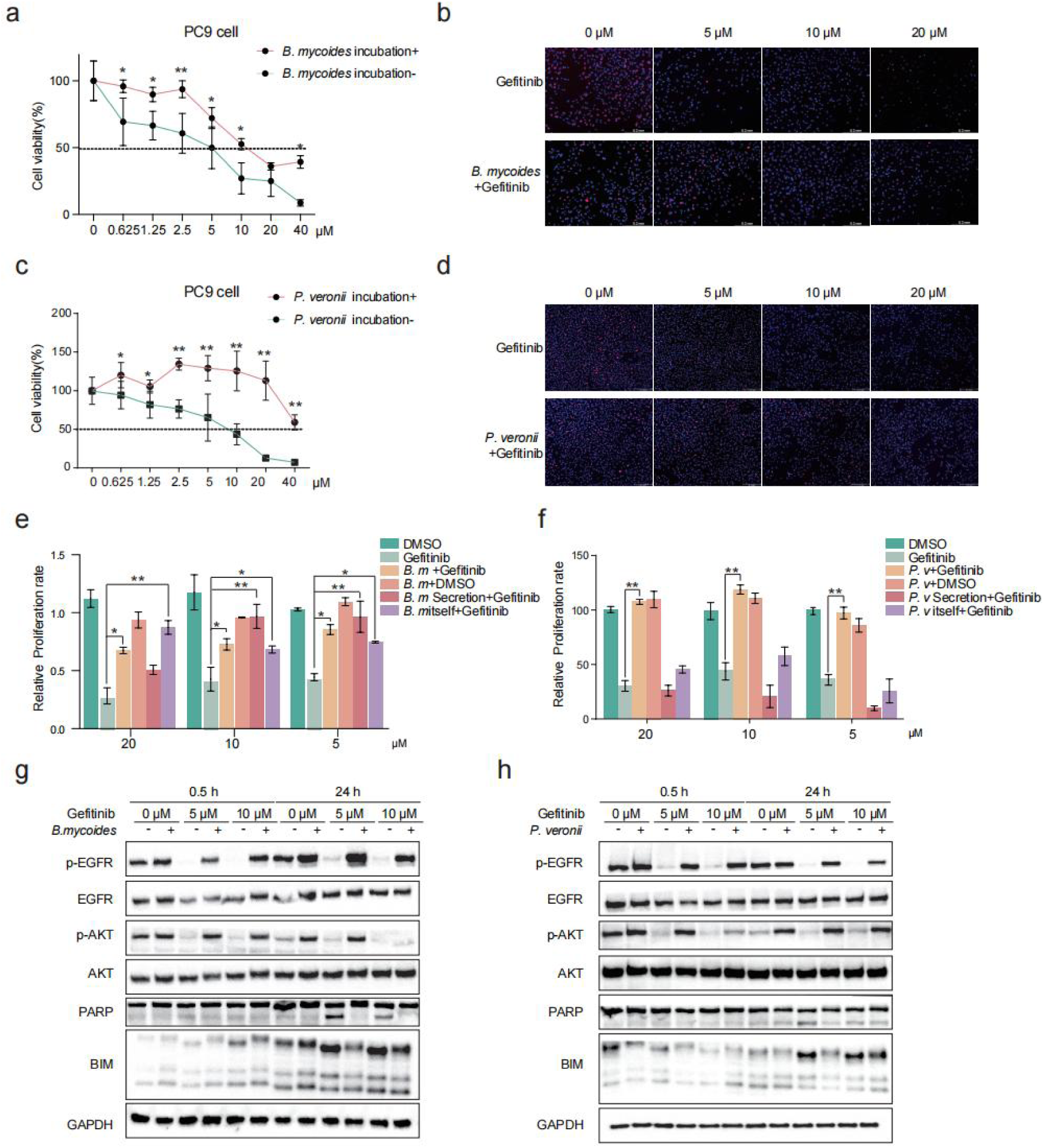
Gefitinib-sensitive cell lines demostrated *B. mycoides* and *P. veronii* could assist cells to tolerate Gefitinib in vitro. **a** Relative proliferation rate of gefitinib-sensitive cell PC9 administered with gefitinib preincubated with/without *B. mycoides*. **b** PC9 cell proliferation was observed under a light microscope in response to gefitinib preincubated with/ without B. *mycoides*. Red flurescenece indicates EdU-positive cells stained with AZD555,blue flurescenece indicates cell nuclei stained with Hoechst 33342. **c** Relative proliferation rate of gefitinib-sensitive cell PC9 administered with gefitinib preincubated with/without *P. veronii*. **d** PC9 cell proliferation was observed under a light microscope in response to gefitinib preincubated with/ without *P. veronii*. Red flurescenece indicates EdU-positive cells stained with AZD555,blue flurescenece indicates cell nuclei stained with Hoechst 33342. **e**,**f**, Relatice proliferation rate of PC9 cells under different treatment conditions. Bacteria, bacterial cells, or bacterial secretions preinbubated with gefitinib at 5μM,10μM,20μM. B. *mycoides* (**e**) and *P. veronii* (**f**). **g**,**h**, Immunoblot analysis showing activity of the EGFR, Akt and PARP cleavage as well as BIM in response to gefitinib (0μM,1μM,10μM), after 0.5h and 24h (+ indicates preincubation with bacteria, − indicates treatment was not given) in PC9 cell lines. GAPDH is the loading control. B. *mycoides* (**g**) and *P. veronii* (**h**).

To delineate whether the observed effects were mediated by bacterial cells or their secreted factors, we implemented a systematic pretreatment protocol for each bacterium: (I) untreated bacterial suspensions containing both cells and their secreted factors; (II) cell-free filtrates obtained through 0.22 μm membrane filtration; and (III) formaldehyde-inactivated bacterial cells. Following pretreatment, these bacterial preparations were subjected to the same gefitinib interaction protocol. For *B. mycoides* (Gram-positive), all three preparations—viable cells, secreted factors, and inactivated cells—demonstrated significant capacity to compromise gefitinib activity (Fig. 2e, Extended Data Fig. 2e). In contrast, for *P. veronii* (Gram-negative), only viable bacterial cells exhibited this inhibitory effect, while cell-free filtrates and inactivated cells showed no significant impact on gefitinib efficacy (Fig. 2f). Further characterization demonstrated differential mechanisms of gefitinib metabolization between the two species. The macroscopic mycelial structure of *B. mycoides* appeared to contribute to its gefitinib-degrading capacity, which was maintained even after formaldehyde inactivation, while *P. veronii* required metabolic activity for gefitinib modification.

To investigate the molecular consequences of bacterial gefitinib modification, we analyzed EGFR signaling dynamics in PC9 cells(Fig. 2g,h). Compared to standard gefitinib treatment, pre-incubated gefitinib resulted in enhanced AKT pathway activation and reduced apoptotic signaling, as indicated by decreased cleavage of poly (ADP-ribose) polymerase (PARP) and altered BIM-EL expression patterns [25–27]. These results collectively demonstrate that bacterial interaction significantly reduces gefitinib’s capacity to inhibit EGFR signaling. Our findings provide compelling evidence that increased intratumoral abundance of *B. mycoides* and *P. veronii* may contribute to the development of gefitinib resistance in NSCLC through distinct bacterial-mediated mechanisms.

To assess the impact of bacterial treatment on the antitumor efficacy of gefitinib in vivo, we established xenograft models using gefitinib-sensitive PC9 cells, as illustrated in Fig. 2a. Tumors treated with gefitinib pre-incubated with either *B. mycoides* or *P. veronii* exhibited a significantly reduced treatment response compared to those treated with gefitinib alone (*B. mycoides*: p=0.038; *P.* veronii: p=0.087) (Fig. 3b-d, e-g). Notably, while gefitinib alone nearly eradicated the tumors, gefitinib pre-incubated with bacteria failed to achieve comparable efficacy. This diminished activity was not attributable to systemic toxicity, as body weights remained stable across all groups (Extended Data Fig. 3a-d). Consistent with the observed tumor growth, immunohistochemical analysis revealed elevated Ki-67 expression in tumors treated with bacterial-metabolized gefitinib (Fig. 3h), suggesting enhanced proliferative capacity. These findings indicate that both *B. mycoides* and *P. veronii* can functionally impair the therapeutic efficacy of gefitinib, potentially promoting tumor cell survival and contributing to drug resistance.

**Fig 3.**
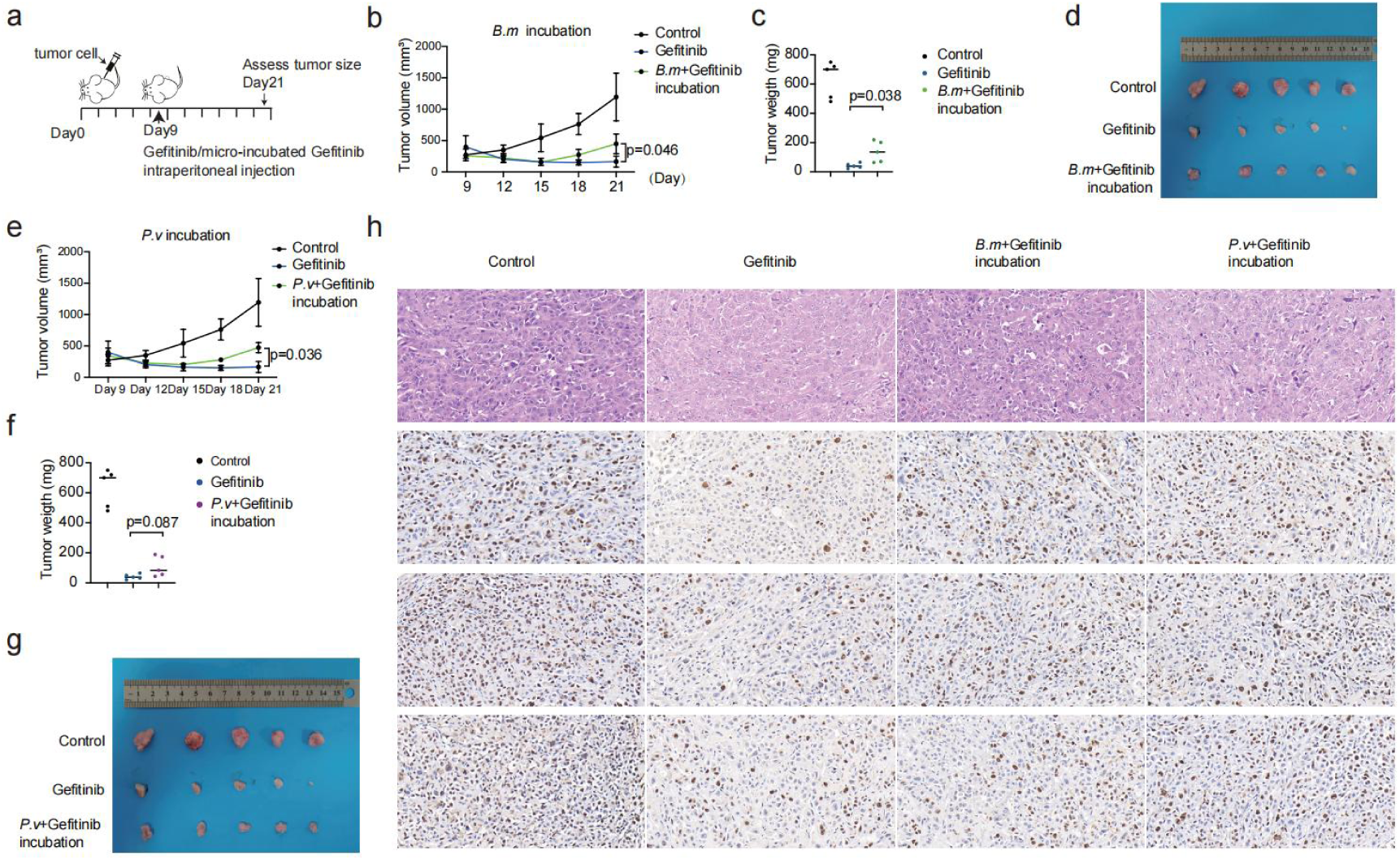
*B. mycoides* and *P. veronii* could assist cells to tolerate Gefitinib in vivo. **a,** Schematic diagram of PC9 xenograft model. Gefitinib and gefitinib preincubate with bacteria was intraperitoneal injected every day. **b,** Tumor volume was measured using calipers to determine the length (L) and width (W) diameters after indicated treatment with *B. mycoides*, and the volume was calculated using the formula: volume= 0.52*(Lx W^2^). **c**, Tumor weight of xenograft model after indicated treatment with *B. mycoides*. **d,** The size of xenograft tumors in mice under different treatment conditions with *B. mycoides*. **e**, Tumor volume was measured using calipers to determine the length and width diameters after indicated treatment with *P. veronii*. **f,** Tumor weight of xenograft model after indicated treatment with *P. veronii*. **g**, The size of xenograft tumors in mice under different treatment conditions with *P. veronii*. **h**, Immunohistochemical stain of tumors stained with Rabbit anti-Ki67 in response to different treatment conditions.

### *B. mycoides* and *P. veronii* metabolize Gefitinib in vitro

To investigate the mechanism underlying *B. mycoides*-induced tumor cell resistance to gefitinib, we co-incubated gefitinib with *B. mycoides* in cell culture medium and analyzed the resulting medium using high-performance liquid chromatography-tandem mass spectrometry (HPLC-MS/MS)(Fig. 4a). The experiments demonstrated a time-dependent decrease in gefitinib concentration (Fig. 4b) . We further investigated the fragmentation patterns of gefitinib and its metabolites, comparing them using CD software. A total of 1,575 metabolites derived from gefitinib were analyzed, including 705 in positive ionization mode and 870 in negative ionization mode (Table S2). After excluding 0-hour interaction products and metabolites undetected in quality control (process shown in Extended Data Fig. 4a), we identified 24 potential metabolites resulting from *B. mycoides*-mediated gefitinib metabolism (Extended Data Fig. 4b). These included 13 in positive ionization mode and 11 in negative ionization mode, with 6 metabolites generated through redox reactions and 18 undergoing subsequent conjugation reactions (Table S3). The temporal profiles of all metabolites are presented. Fig. 4c highlighting primary redox-generated metabolites. The metabolites obtained through conjugation reactions are presented in Fig. 4d-e.

**Fig 4.**
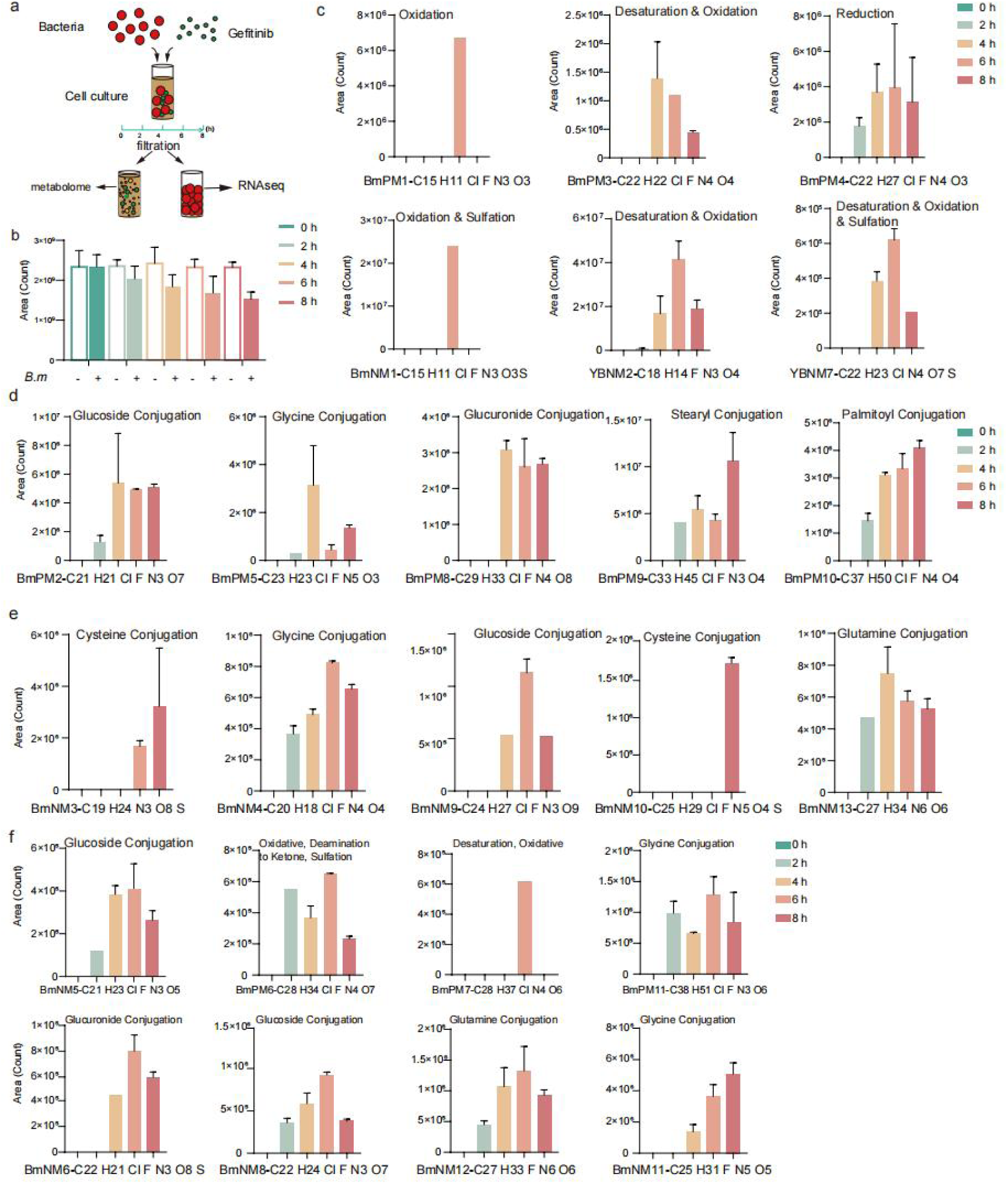
*B. mycoides* exhibits the ability to metabolize gefitinib in vitro. **a,** Schematic diagram of the incubation between *B. mycoides* and gefitinib in vitro. **b,** Time-dependent decrease in gefitinib concentration incubated with *B. mycoides*. **c,** Time-course analysis of six potential gefitinib metabolites generated through redox metabolism by *B. mycoides* using LC-MS. **d,e,** Time-dependent profiles of potential gefitinib conjugative metabolites generated by *B. mycoides* using LC-MS in the positive ionization mode (**d**) and the negative ionization mode (**e**). **f**, Time-dependent profiles of the remaining potential gefitinib conjugative metabolites generated by *B. mycoides* using LC-MS.

Using the secondary fragment structures of metabolites and CD software’s FISH function, we analyzed the structural formulas of the 24 metabolites. By referencing the MS2 spectrum of gefitinib, we predicted metabolite structures and validated them against the MS2 spectra of the predicted metabolites (Extended Data Fig. 4c-d and Extended Data Fig. 5 (negative mode), Extended Data Fig. 4e-f and Extended Data Fig. 6 (positive mode)). As we known, the main pathways of drug metabolism include: phase I reactions, such as oxidation, reduction, and hydrolysis reactions; and phase II reactions, which are conjugation reactions [28]. Our analysis revealed three primary metabolic transformations: oxidation (25%, 6/24), reduction (4.2%, 1/24), and conjugation (70.8%, 17/24) (Table 3). Conjugation substrates were predominantly amino acids or sugar molecules, with oxidation or reduction typically preceding conjugation reactions.

Notably, during the process of predicting metabolite structures, we also uncovered a sequential transformation pathway involving multiple enzymatic modifications. Specifically, we observed that Gefitinib undergoes oxidative metabolism to form BmPM1 (Extended Data Fig. 5e), which is subsequently converted to BmNM1 through sulfidation (Extended Data Fig. 5c). Further metabolic processing of BmNM1 via deamination leads to the formation of the ketone derivative BmNM6. This metabolic cascade is supported by the temporal detection pattern of these metabolites: while both BmPM1 and BmNM1 were first detected at 6 h (Fig. 4c), BmNM6, the downstream metabolite of BmNM1, was detectable as early as 4 h and reached its peak concentration at 6 h (Fig. 4f). This temporal pattern suggests that BmPM1 and BmNM1 were likely synthesized by 4 h but were rapidly metabolized to form BmNM6, explaining their absence at the 4 h time point. Based on these observations, we propose that *B. mycoides* initially metabolizes Gefitinib through oxidoreductase-mediated reactions, followed by secondary modifications involving amino acid transferases or glycosyltransferases. This sequential metabolic processing ultimately yields more complex molecular structures with enhanced solubility properties [29].

To elucidate the molecular mechanisms underlying *B. mycoides*-mediated Gefitinib reduction, we performed RNA sequencing to analyze bacterial gene expression profiles (Table 4). Comparative transcriptomic analysis revealed significant alterations in sugar metabolism, intracellular/extracellular transport systems, and charge conduction pathways between Gefitinib-treated bacteria and untreated controls (0 h) (Extended Data Fig. 7a, b). Transcriptional profiling identified 27 significantly upregulated genes (log(FC) > 2, where FC represents fold change relative to 0 h, padj<0.01; Table 5), including several genes encoding redox-active proteins (Extended Data Fig. 7c). These redox-related genes may facilitate the metabolic transformation of Gefitinib through oxidation-reduction reactions. Furthermore, we identified 4 genes associated with starch and sucrose metabolism, which are closely linked to bacterial biotransformation (Extended Data Fig. 7d). And, 10 differentially expressed genes associated with bacterial transport systems, 5 of which exhibited remarkable upregulation exceeding 100-fold (Extended Data Fig. 7e). This substantial induction of transport-related genes likely supports the intracellular accumulation of metabolic precursors required for the formation of covalent macromolecular complexes. The remaining upregulated genes were predominantly associated with stress response pathways, including multiple transcriptional regulators (Extended Data Fig. 7f). These regulatory elements may orchestrate the coordinated expression of metabolic enzymes involved in Gefitinib biotransformation.

Transcriptional and metabolic analyses were also conducted following the incubation of *P. veronii* with gefitinib. Consistent with the observations in *B. mycoides*, the concentration of the parent drug, gefitinib, decreased in a time-dependent manner (Extended Data Fig. 7g). However, comparative analysis identified only 11 potential metabolites, with metabolic pathways primarily involving redox and conjugation reactions (Table 6). The structures of all metabolites were analyzed using CD software, and the results are presented in Extended Data Fig. 8. Notably, the structures of five metabolites remained unresolved. Transcriptomic analysis of *P. veronii* was performed using the same criteria as for *B. mycoides* (Table 7). Given that the *P. veronii* transcriptomic dataset comprised four replicates — more stringent than the three replicates used for *B. mycoides* — fewer differentially expressed genes met the screening thresholds (27 in *P. veronii* compared to 148 in *B. mycoides*). Although not upregulated at all time points as observed in *B. mycoides*, these genes responded to gefitinib metabolism during the initial stages of *P. veronii*-gefitinib interaction. Furthermore, genes associated with redox processes and sulfur-iron transport exhibited significant upregulation in response to gefitinib metabolism (Extended Data Fig. 7h,i). All significantly upregulated genes in *P. veronii* linked to gefitinib metabolism were categorized into four functional groups: stress regulation, redox processes, substance transport, and other functions of unknown significance (Extended Data Fig. 7j-m, Table 7). These findings suggest that *P. veronii* employs a distinct yet partially overlapping set of metabolic and transcriptional responses compared to *B. mycoides*, highlighting species-specific adaptations to gefitinib exposure. This study provides new insights into the microbial degradation of gefitinib and underscores the potential role of environmental microbes in the biotransformation of pharmaceutical compounds.

### *B. mycoides* and *P. veronii* metabolize Gefitinib in vivo

To investigate whether the observed metabolism of gefitinib could be attributed to *B. mycoides* or *P. veronii* in vivo, we modified the treatment protocol as illustrated in Fig. 5a. The animal model was established as previously described, with adjustments made to the interaction pattern between the bacteria and gefitinib. Following peritumoral injection of *B. mycoides* or *P. veronii*, tumor growth in PC9-bearing mice was not reduced compared to gefitinib treatment alone. This suggests that the presence of *B. mycoides* and *P. veronii* significantly impairs the therapeutic efficacy of gefitinib (*B. mycoides*, Fig. 5b-d; p value= 0.0013; *P. veronii*, Fig. 5e-g; p value= 0.013). Ki-67 expression was also assessed and found to be elevated in the groups receiving peritumoral bacterial injection (Fig. 5h) compared to the PBS control group. In contrast, no significant change in Ki-67 expression was observed in groups treated with peritumoral bacterial injection alone (without gefitinib; Extended Data Fig. 9e). Furthermore, tumor tissues were collected for metabolite analysis (Table 8). Notably, the metabolite B.mPM9-C33 H45 Cl F N3 O4, which is produced by *B. mycoides* during gefitinib metabolism in vitro, was exclusively detected in tumor tissues from mice receiving peritumoral *B. mycoides* injection. Similarly, the metabolite P.vPM2-C24 H29 Cl N4 O3, generated by *P. veronii* during gefitinib metabolism in vitro, was only identified in tumor tissues from mice receiving peritumoral *P. veronii* injection. These findings provide direct evidence that *B. mycoides* and *P. veronii* metabolize gefitinib in vivo, potentially contributing to the observed reduction in therapeutic efficacy. These results demonstrate that both *B. mycoides* and *P. veronii* are capable of metabolizing gefitinib in vivo, leading to the production of distinct metabolites and a consequent reduction in the drug’s antitumor efficacy. This highlights the potential impact of microbial activity on the pharmacokinetics and therapeutic outcomes of anticancer drugs, underscoring the need to consider microbial interactions in cancer treatment strategies.

**Fig 5.**
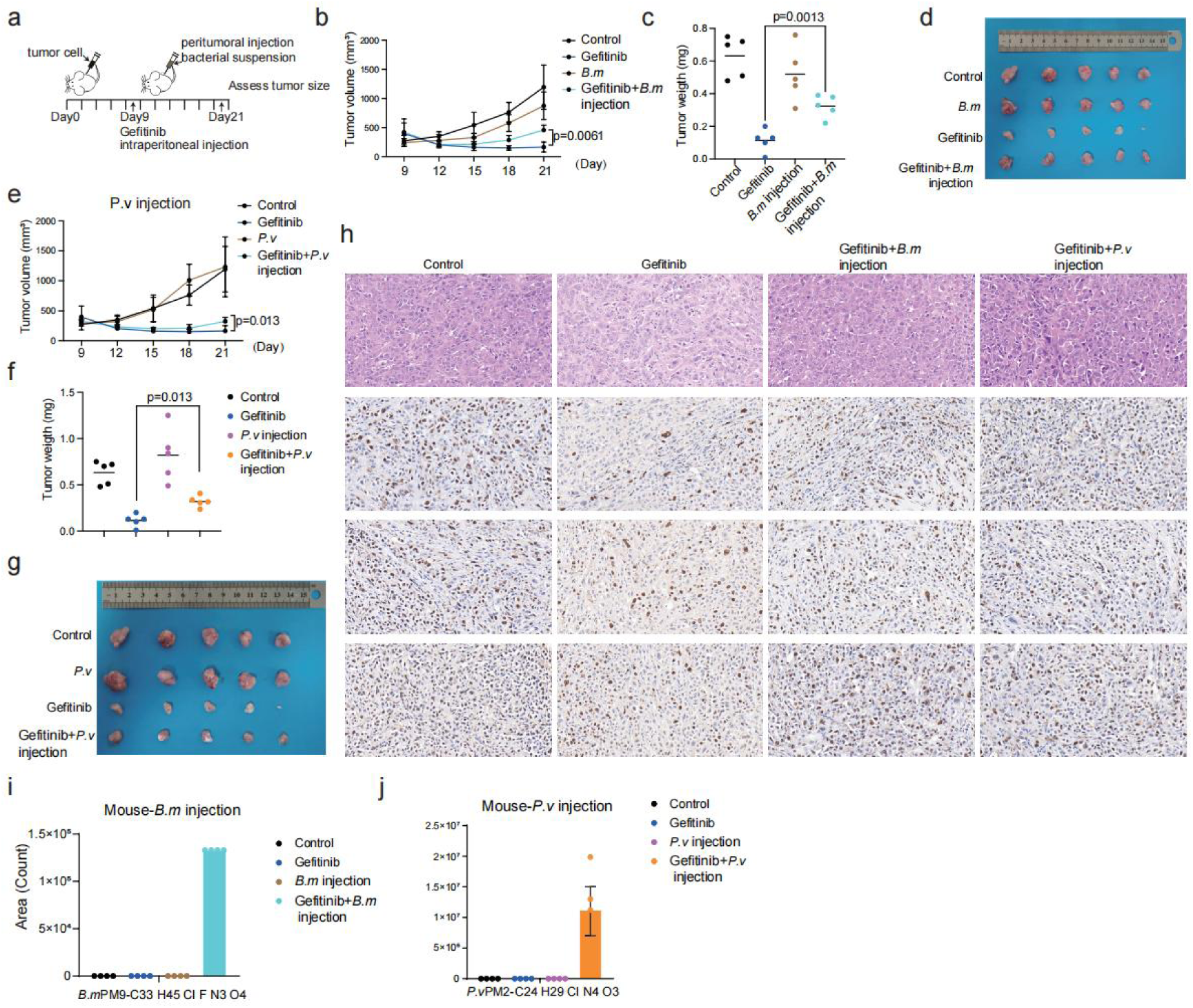
*B. mycoides* and *P. veronii* alleviated gefitinib‘s inhibtion by metabolism in vivo. **a,** Schematic diagram of PC9 xenograft model. Gefitinib and gefitinib preincubate with bacteria was intraperitoneal injected every day while bacteria was pertumoral injected once every three days . **b**, Tumor volume was measured using calipers to determine the length and width diameters after injection with *B. mycoides* and gefitinib, and the volume was calculated using the formula: volume= (Lx W2)/2. **c**, Tumor weight of xenograft model after injection with *B. mycoides* and gefitinib. **d**, The size of xenograft tumors in mice under injection treatment with *B. mycoides*. **e**, Tumor volume was measured using calipers to determine the length and width diameters after injection with *P. veronii*. **f**, Tumor weight of xenograft model after injection with *P. veronii*. **g**, The size of xenograft tumors in mice under injection treatment with *P. veronii*. **h**, Immunohistochemical stain of tumors stained with Rabbit anti-Ki67 after injection with gefitinib and bacterias. **i, j,** Metabolites of gefitinib was found in mice tumors injected with *B. mycoides* (**i**) and *P. veronii* (**j**).

**Fig 6.**
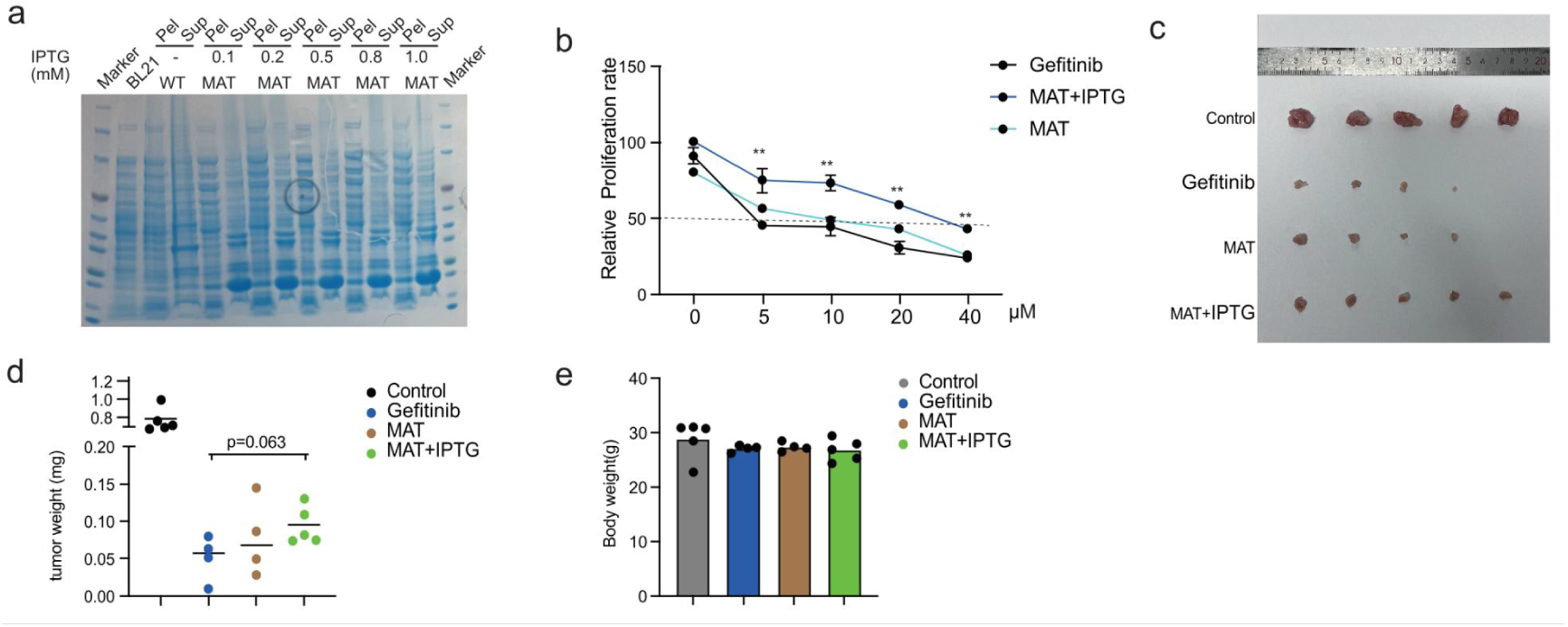
The expression of maltose acetyltransferase enables *E.coli* to metabolize gefitinib. **a**, The results of polyacrylamide gel stained with Coomassie Brilliant Blue. After electrophoresis, the protein bands of samples under different induction conditions are clearly visible in this figure. *BL21* represents the competent cells, MAT(*Bm-RS09505*) represents maltose acetyltransfease, Sup represents the supernatant after sonication and centrifugation, and Pel represents the pellet after sonication and centrifugation. **b**, Relative proliferation rate of PC9 in response to gefitinib preincubated with *E.coli* with or without MAT expression. **c**, The size of xenograft tumors in mice under different treatment conditions with *E.coli* with or without MAT expression. **d**, Tumor weight of xenograft in response to indicated treatment with *E.coli.* **e**, The body weight of mice in response to indicated treatment with *E.coli*.

### Escherichia coli expressing maltose acetyltransferase has acquired the ability to metabolize gefitinib

To investigate which bacterial proteins are involved in the metabolism of gefitinib, we screened for differentially expressed genes in bacteria under gefitinib stimulation. Transcriptomic data analysis revealed that, in response to drug stimulation, the expression of multiple redox proteins and components of the bacterial two-component system were significantly upregulated, including flavodoxin, TIpA disulfide reductase, α,α-trehalose phosphorylase, and maltose acetyltransferase. To validate the function of these proteins, we heterologously expressed these genes in *Escherichia coli*, a strain that does not participate in drug metabolism and applied the same pre-incubation treatment protocol, followed by assessing the inhibitory efficacy of gefitinib on cells. The results showed that pre-incubation of gefitinib with *E. coli* expressing maltose acetyltransferase significantly attenuated the drug’s inhibitory effect on cells, whereas similar phenomenon was not observed in other protein expression groups. This finding suggests that maltose acetyltransferase may play a critical role in the metabolism of gefitinib by *B. mycoides*.

To further validate this discovery, we conducted similar experiments in a mouse xenograft tumor model and obtained results consistent with those from the in vitro experiments. These results indicate that under gefitinib stimulation, *B. mycoides* upregulates the expression of maltose acetyltransferase, metabolizing gefitinib into inactive forms, thereby reducing the effective concentration of gefitinib in cells and ultimately promoting drug resistance. However, the specific molecular mechanism by which maltose acetyltransferase metabolizes gefitinib remains unclear. Future research should focus on elucidating the enzymatic pathways involved, including potential intermediate metabolites and regulatory networks, to provide new targets and strategies for overcoming tumor drug resistance.

## Discussion

The role of tumor-resident microorganisms in mediating resistance to targeted therapies in lung cancer remains poorly characterized. Here, through an eight-year longitudinal study, we established a unique cohort of matched clinical samples collected from NSCLC patients before and after the development of drug resistance. Using third-generation long-read sequencing technology, we identified significant enrichment of two bacterial species, *B. mycoides* and *P. veronii*, within tumor tissues of treatment-resistant patients. While prior studies have extensively explored the role of oral and gut microbiota in modulating colorectal cancer immunotherapy [30–33], our findings uncover a previously unrecognized function of tumor-resident microbiota in driving targeted therapy resistance in lung cancer. This discovery provides critical insights into the clinical management of NSCLC patients receiving targeted therapies. Mechanistic studies reveal that *B. mycoides* and *P. veronii* may directly influence therapeutic outcomes by metabolizing gefitinib, both in vivo and in vitro. Specifically, we identified two metabolites, B.mPM9-C33 H45 Cl F N3 O4 and P.vPM2-C24 H29 Cl N4 O3, generated through the bacterial metabolism of gefitinib. These bacterial species and their metabolic byproducts represent potential non-invasive biomarkers for predicting response to targeted therapies. Furthermore, our data suggest that targeting tumor-resident *B. mycoides* and *P. veronii* populations could serve as a novel therapeutic strategy to improve treatment efficacy in NSCLC, akin to microbiota-based interventions currently under development for colorectal cancer immunotherapy [34].

Diverging from prior research that primarily examined host-driven adaptations in reaction to targeted drug resistance [35, 36], this study elucidate the metabolic pathways and bacterial active substances through which tumor-resident microbes enable tumor cells to tolerate EGFR-TKI drugs and metabolize gefitinib. These results significantly advance our understanding of acquired resistance to EGFR-TKI targeted therapies in patients with NSCLC. By uncovering the mechanisms by which tumor-resident microbes influence targeted drug resistance, our study provides new insights for overcoming acquired resistance and highlights the importance of considering microbial interactions in cancer treatment strategies. These findings also raise broader questions about the role of tumor-resident microbes in modulating the pharmacokinetics and acquired resistance of targeted anticancer drugs, as well as the downstream effects of drug-induced changes in microbial community structure and function on drug efficacy and toxicity. Although our in vitro and in vivo experiments support the hypothesis that *B. mycoides* and *P. veronii* in lung tumor tissues impair the efficacy of EGFR-TKI drugs, further studies are needed to evaluate the broader impact of this metabolic transformation on lung tissue and potential side effects. Regrettably, despite repeated attempts, we were unable to isolate clinical strains of these bacteria, which remains a limitation of this study. Future studies should prioritize the isolation and characterization of clinical strains, as these would provide more robust evidence and may uncover additional mechanisms by which intratumoral bacteria modulate drug efficacy.

Previous studies have demonstrated that intratumoral microbes can modulate tumor cell sensitivity to therapies through diverse mechanisms, thereby contributing to treatment resistance. *Leore T. Geller et al.* reported that in pancreatic ductal adenocarcinoma (PDAC), *M. hyorhinis* metabolizes the chemotherapeutic agent gemcitabine into inactive metabolites via cytidine deaminase, leading to chemoresistance [7]. Similarly, *Ni Wang* et al. revealed that in colorectal cancer, *Fusobacterium nucleatum* activates the Hippo signaling pathway and upregulates BCL2 expression, thereby suppressing chemotherapy-induced pyroptosis and promoting chemoresistance [37]. These findings underscore the ability of intratumoral microbes to enhance tumor cell resistance by altering cellular signaling and cell death pathways. In contrast to drug resistance driven by EGFR secondary mutations or bypass signaling activation [38–40], we investigated a novel mechanism by which tumor-resident microbiota confer resistance to targeted therapies. Specifically, we observed that in response to gefitinib treatment, functional genes were upregulated in both *B. mycoides* and *P. veronii*, yet distinct metabolic outcomes were observed. This divergence may be attributed to their differing bacterial classifications. As a Gram-positive bacterium, *B. mycoides* possesses an extensive repertoire of cell wall-associated enzymes and transport proteins, which may facilitate the intracellular retention and metabolism of gefitinib [41]. In contrast, *P. veronii*, a Gram-negative bacterium, likely expels metabolites through outer membrane porins and efflux systems [42]. This mechanistic distinction may explain why *P. veronii* generated fewer metabolites upon gefitinib exposure compared to *B. mycoides*.

Moreover, under stress conditions, *B. mycoides* exhibits higher gene expression diversity and metabolic flexibility, capable of metabolizing gefitinib through redox reactions and conjugation reactions. In contrast, *P. veronii* may primarily generate a narrower range of metabolites via redox reaction [41]. It is worth noting that the genome of *B. mycoides* may be more inclined to acquire new metabolic capabilities through horizontal gene transfer [43]. While this study has primarily focused on the direct effect of *B. mycoides* and *P. veronii* on drug metabolism, previous research by Lauren et al. has shown that Lactobacillus iners could induce chemoresistance in colorectal cancer through metabolic rewiring. Lauren also pointed out that *L. iners* has also been identified in lung, head and neck, and skin cancers, and this observation was consistent with our results. *L. iners* in NSCLC was strongly associated with decreased RFS, which suggested a potential role in drug resistance in NSCLC [6]. Studies have shown that the decrease in pH could reduce the uptake of gefitinib by tumor cells [44]. In paired samples, we also observed an enrichment of *L. iners* in post-resistance patient samples, but this result was not further validated in unpaired samples, warranting more validation and experiments.

Clinically, for NSCLC patients with EGFR mutation-positive status, especially those newly diagnosed and treatment-naïve, first-generation EGFR-TKIs such as gefitinib are often the initial therapeutic choice[45]. However, for patients with EGFR T790M mutations or brain metastases, third-generation EGFR-TKIs are typically administered directly [46]. In this study, we included 13 non-paired samples from patients who developed resistance to third-generation EGFR-TKIs. Among these samples, *B. mycoides* was detected in 2 out of 13 cases, while *P. veronii* was found in 5 out of 13 cases (Table 9). This suggests that the bacteria identified in our study, which are capable of metabolizing first-generation drugs like gefitinib, may also be present in patients resistant to third-generation EGFR-TKIs. However, whether these bacteria can metabolize third-generation drugs remains unclear and warrants further investigation with larger sample sizes and mechanistic studies to elucidate their potential role in resistance to newer-generation therapies.

From first-generation drugs like gefitinib and erlotinib to third-generation drugs such as osimertinib and rociletinib, significant clinical benefits have been achieved in the treatment of NSCLC with EGFR mutations. However, the inevitable emergence of drug resistance remains a major challenge. This study provides new insights into how tumor-resident microbiota facilitate the development of resistance in tumor cells. By identifying microbes enriched in tumors after gefitinib resistance, our findings supported the “survival of the fittest” hypothesis. We hypothesized that similar variations in microbial composition, abundance, and enrichment patterns may occur in tumor tissues resistant to other TKI drugs. These enriched microbes may induce drug resistance through mechanisms such as metabolizing pharmacodynamic active components of drugs, releasing specific metabolites, or affecting cellular autophagy. Tumor microbiota play a crucial role in regulating tumor progression and treatment outcomes. Harnessing the microbiota could serve as a novel therapeutic strategy for cancer diagnosis, prognosis, and treatment. Although current research has not fully clarified whether changes in the tumor microbiota are a cause or consequence of cancer development, future studies will continue to explore these complex relationships to develop more effective treatment strategies. Despite the limited sample size in this study, it still provides a novel potential mechanism by which microbiota facilitate drug resistance in tumor cells. In future research, more samples need to be enrolled to determine whether the abundance of specific microbes is associated with poor prognosis or the emergence of resistance. In clinical treatment, the combination of antibiotics and targeted drugs could offer a new therapeutic approach for NSCLC and other cancers.

## Methods

### Human samples collection

Between 2015 and 2023, 128 tumor samples were collected from patients with NSCLC after obtaining informed consent. These samples were divided into a pilot cohort (14 paired tumor samples before and after EGFR-TKI resistance) and a validation cohort (114 unpaired tumor samples before and after EGFR-TKI resistance). Clinical information was obtained from the electronic medical record and is provided in Supplementary Table S1. Lung tumor tissues were collected from patients who underwent lung cancer resection, percutaneous lung biopsy, or bronchoscopic biopsy at West China Hospital, Sichuan University. All samples were confirmed to be tumor tissue based on pathological assessment. DNA was extracted from these samples for bacterial quantification and 16S rRNA library preparation.

### Bacteria culture

B. *mycoides* (bio-120970) *P. veronii* (bio-02408) and Lactobacillus iners(bio-107295) were purchased from Biobw. *B. mycoides* were cultured in Tryptic Soy Agar plate at 30 °C aerobically, *P. veronii* were cultured in PVG plate(peptone 10 g, yeast extract 5g, glucose 1g, agar 15g,distilled water 1L) at 25°C aerobically, *Lactobacillus iners* were cultured in an anaerobic chamber Hypoxystation in Tryptic Soy Agar plate +5% sheep blood (Solarbio-#TX0030). The bacteria were passaged every day or once every two days.

### Cell culture

PC-9,H1975 and A549 were obtained from ATCC.H1975 and A549 cells were cultured in DMEM supplemented with 10% FBS and 100 U ml–1 penicillin-streptomycin. PC-9 cells were cultured in RPMI 1640 supplemented with 10% FBS and 100 U ml–1 penicillin-streptomycin. All cells were incubated in a humidified cell culture incubator with 5% CO2 at 37°C. Cells were used for no longer than 12months before being replaced and were routinely tested for Mycoplasma to ensure the accuracy of experimental data.

### Cell proliferation assay

PC-9 cells were seeded in 96-well plate at a densitity of 2000 cells per well in a total of 100 μl culture medium and incubated at 37℃ 5% CO2. Bacteria were per-incubated with gefitinib for 4 hours and then filtered out. Once the cells had adhered to the bottom of the well,the medium was replaced with fresh medium containing varying concentrations of gefitinib, or medium that had been pre-incubated with bacteria. The cells were cultured for an additional 48hours. Subsequently, 10μl cck-8 reagent was added to each well, and the aborbance was measured at a wavelength of 450nm. Cell relative proliferation rate(%)=[(administration group A-Negative control group A)/(control group A-Negative control groupA)]*100%.The half-maximal inhibitory concentration (IC50) was calculated based on the relative proliferation curve using GraphPad Prism v9.0.

### EdU assay

PC-9 cells were seeded in 96-well plate at a density of 2000 cells per well in a total of 100 μl culture medium and incubated at 37℃, 5% CO2.Bacteria were per-incubated with gefitinib for 4 hours and then filtered out. Once the cells had adhered to the bottom of the well,the medium was replaced with fresh medium containing varying concentrations of gefitinib, or medium that had been pre-incubated with bacteria. The cells were cultured for an additional 48hours. Add a final concentration of 10 μM Edu working solution to each well and culture for an additional 2hours after EdU labeling of cells was completed, remove the culture medium and add the fix solution, then fix at room temperature for 15mins, After removing the fix solution, wash the cells with washing solution three times for 5 minutes each. Then, permeabilize the cells with permeabilization solution for 15 minutes at room temperature. Finally, wash the cells again with washing solution twice for 5 minutes each. Remove the washing solution and add the Click reaction solution. Incubate at room temperature protected from light for 30minutes. wash the cells with washing solution three times for 5 minutes each. For cell nucleus labeling, Hoechst 33342 was added and incubated in the dark at room temperature for 10 minutes. Fluorescence detection was conducted using a Leica DMi8 fluorescence microscope, with measurements taken at a wavelength of 555nm for Azide and 346nm for Hoechst, respectively. Cell counting is performed by Imaris software after separating the two fluorescence signal channels.

### Mouse Xenograft studies

Male BALB/c nude mice aged 6 weeks were purchased from Nanjing Gempharmatech Co., Ltd. (Nanjing, China), and were acclimatized for 1 week. 2×10^6^ PC-9 tumor cell within 50% Matrigel^®^(coring) were subcutaneously inoculated into the right flanks of the mice. After about 9 days, gefitinib with/without pre-incubation with bacteria was administered at the dose of 12.5mg/kg every day by intraperitoneal injection. Similarly, 5×10^6^ CFU bacteria were injected into tumor once every three day. Fifteen days after first injection, mice were sacrificed to characterize tumor weight and possible metabolites.

### Immunohistochemistry

The subcutaneous tumor tissue harvested from nude mice was immediately fixed in 10% neutral buffered formalin (NBF) for 24–48 hours, followed by dehydration through graded ethanol solutions (70%, 80%, 95%, and 100%), xylene clearing, and paraffin embedding. Tissue sections, 4–5 μm thick, were prepared using a microtome, mounted on glass slides, and dried in a 60℃ oven for 1 hour. The sections were deparaffinized with xylene and rehydrated through graded ethanol solutions, then rinsed with distilled water. Antigen retrieval was performed in 0.01 M sodium citrate buffer (pH 6.0) by microwave heating for 15 minutes, followed by cooling to room temperature and washing with distilled water. Endogenous peroxidase activity was blocked with 3% hydrogen peroxide at room temperature for 10 minutes, and non-specific binding sites were blocked with 5% bovine serum albumin (BSA) or 10% normal goat serum for 30 minutes. Sections were incubated with a rabbit anti-Ki67 monoclonal antibody at the dilution of 1:200 overnight at 4℃ in a humidified chamber, followed by washing with PBS and incubation with an HRP-conjugated secondary antibody for 30 minutes at room temperature. After rinsing with PBS, DAB working solution was applied for chromogenic detection, and the reaction was monitored under a microscope for 3–5 minutes before termination with distilled water. Hematoxylin was used for counterstaining the nuclei, followed by thorough rinsing, dehydration through graded ethanol, xylene clearing, and mounting with neutral resin. Once the mounting medium solidified, the sections were examined under a light microscope to evaluate Ki67 staining.

### Western Blotting

The total protein of the Gefitinib-treated PC-9 and the Gefitinib-Bacteria-coincubation treated PC-9 cells were lysed using Radioimmonoprecipitatipn assay(RIPA)buffer (Beyotime) containing a protease inhibitor cocktail(MedChemExpress).Samples were collected and centrifuged at 14,000 r.p.m. for 20 min at 4 ℃. Protein concentration were quantified by bicinchoninic acid (BCA) assay. Equal amounts of protein(40μg) were loaded onto SDS-polyacrylamide gel electrophoresis gels, transferred to a polyvinylidene difluoride membrane(Millipore) and incubated with the indicated primary antibodies. Proteins were detected via incubation with horseradish peroxidase-conjugated secondary antibodies, ImmobilonR Western Chemiluminescent HRP Substrate(Millipore). Antibodies for EFGR phospho-Akt (S473), Akt, cleaved PARP, BIM, phospho-BIM (S69), GAPDH were purchased from Cell Signaling Technology; phospho-EGFR (Y1068) was purchased from Abcam;

#### In Vitro Protein induction

The expression plasmid was transformed into *BL21* (DE3) competent cells and cultured on plates containing kanamycin resistance. Single colonies were selected and cultured for sequence verification. Positive single colonies were preserved and subjected to expression detection. A single colony was inoculated into 5 mL of LB medium containing kanamycin resistance and cultured at 37°C with constant shaking for 10-12 hours. The bacterial culture was then inoculated into fresh LB medium containing kanamycin resistance at a ratio of 1:100. The culture was grown at 37°C with constant shaking until it reached the logarithmic growth phase (OD600 approximately 0.6-0.8), at which point IPTG was added to a final concentration of 0.1 mM to 1 mM. The culture was then incubated at 16°C with shaking at 120 rpm for 16 hours.The samples were centrifuged at 12000 rpm at 4°C for 10 minutes. The supernatant was discarded, and the pellet was resuspended in 500 μL of Lysis buffer for sonication. The sonication was performed at a power setting of 100 W, with 3 seconds of sonication followed by 5 seconds of rest, repeated for a total of 3 minutes. After sonication, the samples were centrifuged at 4°C at 12000 rpm for 30 minutes. The supernatant and pellet were collected separately, with the pellet being resuspended in an equal volume of Lysis buffer.The supernatant and pellet proteins were mixed with 5× SDS-PAGE Loading Buffer and boiled for 5 minutes to denature the proteins. The samples were subjected to SDS-PAGE. The gel was washed and stained with Coomassie Brilliant Blue for 1 hour, followed by destaining and imaging.

### Transcriptomics and metabolomics sample preparation

To simulate the conditions in cell lines, 1×10^7^ CFU bacteria were incubated with 40 µM Gefitinib in 1 ml of culture medium. At 0 hour, the mixture was centrifuged at 3000 rpm for 10 minutes at 4°C, and the entire supernatant was collected and snap-frozen in liquid nitrogen for 15 minutes. The bacterial pellet was then quickly washed 3 times with pre-chilled PBS, followed by centrifugation at 4°C and 5000 rpm for 5 minutes after each wash; the supernatant was completely discarded, and the bacterial pellet was collected in a 1.5 ml cryogenic tubes and snap-frozen in liquid nitrogen for 15 minutes. To simulate the physiological conditions of gefitinib within the human body incubate the identical aliquots mixture in a 37°C incubator. The aforementioned procedure was repeated at 2 hours, 4 hours, 6 hours, and 8 hours time point to collect the bacterial pellet and culture medium. The supernatant of the culture medium was prepared for metabolomics sequencing, and the bacterial pellet was prepared for transcriptomics sequencing.The experiment was performed four times to represent biologically independent samples.

### 16S Pacbio sequencing

Total genome DNA from samples was extracted using CTAB/SDS method. DNA concentration and purity was monitored on 1% agarose gels. According to the concentration, DNA was diluted to 1 ng/μL using sterile water.16S rRNA genes of distinct regions were amplified used specific primer with the barcode. All PCR reactions were carried out with TransStart® FastPfu DNA Polymerase (TransGen Biotech).Mix same volume of 1X loading buffer (contained SYB green) with PCR products and operate electrophoresis on 2% agarose gel for detection. PCR products was mixed in equidensity ratios. Then, mixture PCR products was purified with QIAquick@ Gel Extraction Kit (QIAGEN)Sequencing libraries were generated using SMRTbellTM Template Prep Kit (PacBio) following manufacturer’s recommendations. The library quality was assessed on the Qubit@ 4.0 Fluorometer (Thermo Scientific) and FEMTO Pulse system. At last, the library was sequenced on the PacBio Sequel platform.

### RNA-sequencing

Bacteria RNA was extracted by RNAprep Pure Cell/Bacteria Kit following manufacturer’s recommendation. Quality control was performed using the Agilent 2100 Bioanalyzer. Total RNA with poly(A) tail was enriched using oligo(dT) magnetic beads, and the enriched RNA was then randomly fragmented. Using the fragmented mRNA as a template and random oligonucleotides as primers, the first strand of cDNA was synthesized in an M-MuLV reverse transcriptase system. Subsequently, the RNA strand was degraded using RNase H, and the second strand of cDNA was synthesized in a DNA polymerase I system. The purified double-stranded cDNA was then subjected to end repair, addition of an A-tail, and ligation of sequencing adapters. The final library was sequenced using the illumina NovaSeq 6000.

### High-performance liquid chromatography-tandem mass spectrometry (HPLC-MS/MS)

HPLC-MS/MS was performed on a Thermo Scientific LCMS system, which includes a Q Exactive™ HF/Q Exactive™ HF-X, a Vanquish UHPLC, and a Hypesil Gold column. Gefitinib ions monitored: 447.1/127.9 in the positive ionization mode, LC method:Column temperature: 40℃, flow rate: 200µl/min, solvent A: 0.1% formic acid,, solvent B: 100% methanol, column: Hypesil Gold column-c18 2.1 x 100mm,1.9µm, The scan range is selected as m/z 100-1500. The settings for the ESI source are as follows: Spray Voltage: 3.5kV; Sheath gas flow rate: 35psi; Aux Gas flow rate: 10L/min; Capillary Temp: 320°C; S-lens RF level: 60; Aux gas heater temp: 350°C; Polarity: positive, negative. The MS/MS second-level scan is a data-dependent scan.

### Data analysis

#### (PACBIO SEQUENCING)

1. **Quality control**

1.1 Data split

Raw sequences were initially processed through the PacBio SMRT portal. Sequences were filtered for a minimum of 3 passes, and a minimum predicted accuracy of 90% (minfullpass = 3, minPredictedAccuacy = 0.9). The predicted accuracy of 90%, which is defined as the threshold below which a CCS is considered as noise. The files generated by the PacBio platform were then used for amplicon size trimming to remove sequences outside the expected amplicon size (minLength 1340 bp, maxLength 1640 bp). The reads was assigned to samples based on their unique barcode and truncated by cutting off the barcode and primer sequence.

1.2 Chimera removal

The reads were compared with the reference database using UCHIME algorithm (UCHIME Algorithm, http://www.drive5.com/usearch/manual/uchime_algo.html) [47] to detect chimera sequences, and then the chimera sequences were removed [48] . Then the Clean Reads finally obtained.

2. OTU cluster and Species annotation

2.1 OTU Production

Sequences analysis were performed by Uparse software (Uparse v7.0.1001 http://drive5.com/uparse/) [49]. Sequences with ≥97% similarity were assigned to the same OTUs. Representative sequence for each OTU was screened for further annotation.

2.2 Species annotation

For each representative sequence, the SSUrRNA Database [50] of Silva Database (https://www.arb-silva.de/) [51] was used based on Mothur algorithmto annotate taxonomic information.

2.3 Phylogenetic relationship Construction

In order to study phylogenetic relationship of different OTUs, and the difference of t he dominant species in different samples(groups), multiple sequence alignment were conducted using the MUSCLE software (Version 3.8.31 http://www.drive5.com/musc le/) [52].

2.4 Data Normalization

OTUs abundance information were normalized using a standard of sequence number corresponding to the sample with the least sequences. Subsequent analysis of alpha diversity and beta diversity were all performed basing on this output normalized data.

3. Alpha Diversity

Alpha diversity is applied in analyzing complexity of species diversity for a sample through indices, including Observed-species, Chao1, Shannon, Simpson, ACE, Good-coverage. All this indices in our samples were calculated with QIIME (Version1.9.1) and displayed with R software (Version 2.15.3). Two indices were selected to identify Community richness: Chao - the Chao1 estimator (http://www.mothur.org/wiki/Chao); ACE - the ACE estimator (http://www.mothur.org/wiki/Ace); Two indices were used to identify Community diversity: Shannon - the Shannon index (http://www.mothur.org/wiki/Shannon); Simpson - the Simpson index (http://www.mothur.org/wiki/Simpson); One indice to characterized Sequencing depth: Coverage - the Good’s coverage (http://www.mothur.org/wiki/Coverage)

1. Beta Diversity

Beta diversity analysis was used to evaluate differences of samples in species complexity, Beta diversity on both weighted and unweighted unifrac were calculated by QIIME software (Version 1.9.1).

Cluster analysis was preceded by principal component analysis (PCA), which was applied to reduce the dimension of the original variables using the FactoMineR package and ggplot2 package in R software (Version 2.15.3).

Principal Coordinate Analysis (PCoA) was performed to get principal coordinates and visualize from complex, multidimensional data. A distance matrix of weighted or unweighted unifrac among samples obtained before was transformed to a new set of orthogonal axes, by which the maximum variation factor is demonstrated by first principal coordinate, and the second maximum one by the second principal coordinate, and so on. PCoA analysis was displayed by WGCNA package, stat packages and ggplot2 package in R software (Version 2.15.3).

Unweighted Pair-group Method with Arithmetic Means (UPGMA) Clustering was performed as a type of hierarchical clustering method to interpret the distance matrix using average linkage and was conducted by QIIME software (Version 1.9.1).

### Data Preprocessing and Metabolite Identification

The raw data files (.raw) were imported into the CD 3.1 search library software for processing. Each metabolite was subjected to simple screening based on retention time and mass-to-charge ratio (m/z). The peak area was then normalized using the first QC sample to enhance the accuracy of identification. Subsequently, peak extraction was performed by setting parameters such as mass deviation (5 ppm), signal intensity deviation (30%), minimum signal intensity, and adduct ions. The peak area was quantified, and target ions were integrated. Molecular formulas were predicted using the molecular ion peaks and fragment ions, and the results were compared with databases such as mzCloud (https://www.mzcloud.org/), mzVault, and Masslist. Background ions were removed using blank samples. The original quantitative results were standardized using the formula: sample original quantitative value / (sample metabolite quantitative value sum / QC1 sample metabolite quantitative value sum) to obtain relative peak areas. Compounds with a CV greater than 30% in the QC samples were removed, resulting in the identification and relative quantification of metabolites. Data processing was conducted on a Linux operating system (CentOS version 6.6) using software such as R and Python, with specific packages and software versions detailed in the results file readme.

### Data Statistical Analysis

Metabolites identified were annotated using the KEGG database (https://www.genome.jp/kegg/pathway.html), HMDB database (https://hmdb.ca/metabolites), and LIPIDMaps database (http://www.lipidmaps.org/).

For multivariate statistical analysis, the metabolomics data processing software metaX [53] was used to transform the data and perform principal component analysis (PCA) and partial least squares discriminant analysis (PLS-DA) to obtain the VIP values for each metabolite. In univariate analysis, t-tests were conducted to calculate the statistical significance (P-value) of each metabolite between two groups and to determine the fold change (FC) values. The default criteria for differential metabolite selection were VIP>1,

P-value<0.05, and FC≥2 or FC≤0.5.

Volcano plots were created using the R package ggplot2, integrating three parameters: VIP value, log2(FoldChange), and −log10(P-value) to select metabolites of interest. Cluster heatmaps were generated using the R package Pheatmap, with z-score normalization applied to the metabolite data.

Correlation analysis between differential metabolites (Pearson correlation coefficient) was performed using the R function cor(), and statistical significance was assessed using the cor.mtest() function in R, with P-value<0.05 considered statistically significant. Correlation plots were created using the corrplot package in R.

Bubble plots were also generated using the R package ggplot2, utilizing the KEGG database to study the function and metabolic pathways of metabolites. A pathway was considered enriched when x/n>y/n, and pathways with P-value<0.05 were deemed significantly enriched.

## Acknowledgements

This project was funded by the National Natural Science Foundation of China (82173182, L.D.), the National Natural Science Foundation of China (32300003, H.W.), the 1·3·5 project for disciplines of excellence, West China Hospital, Sichuan University (ZYYC23024, D.X.), Science and Technology Program of Sichuan (2023NSFSC1939, L.D.).

## Author contributions

D.X. and H.W. conceptualized the project. H.W., J.R.Z., J.X.Z. and W. H. developed the methodology. H.W., J.R.Z., J.X.Z., W. H., and P.W.T. conducted investigations. P.W.T., W.M.L., D.L., and D.X. acquired resources. H.W., J.R.Z., J.X.Z., W. H., J.Z., L.S., M.L.L., X.Y.L. and Y.Q.L. performed visualization. D.X., D.L. and H.W. acquired funding. D.X. and D.L. administered and supervised the project. H.W.J.R.Z., J.X.Z. and W. H. wrote the original draft. D.X. and D.L. reviewed and edited the manuscript.

---

**Extended Data Fig. 1.**
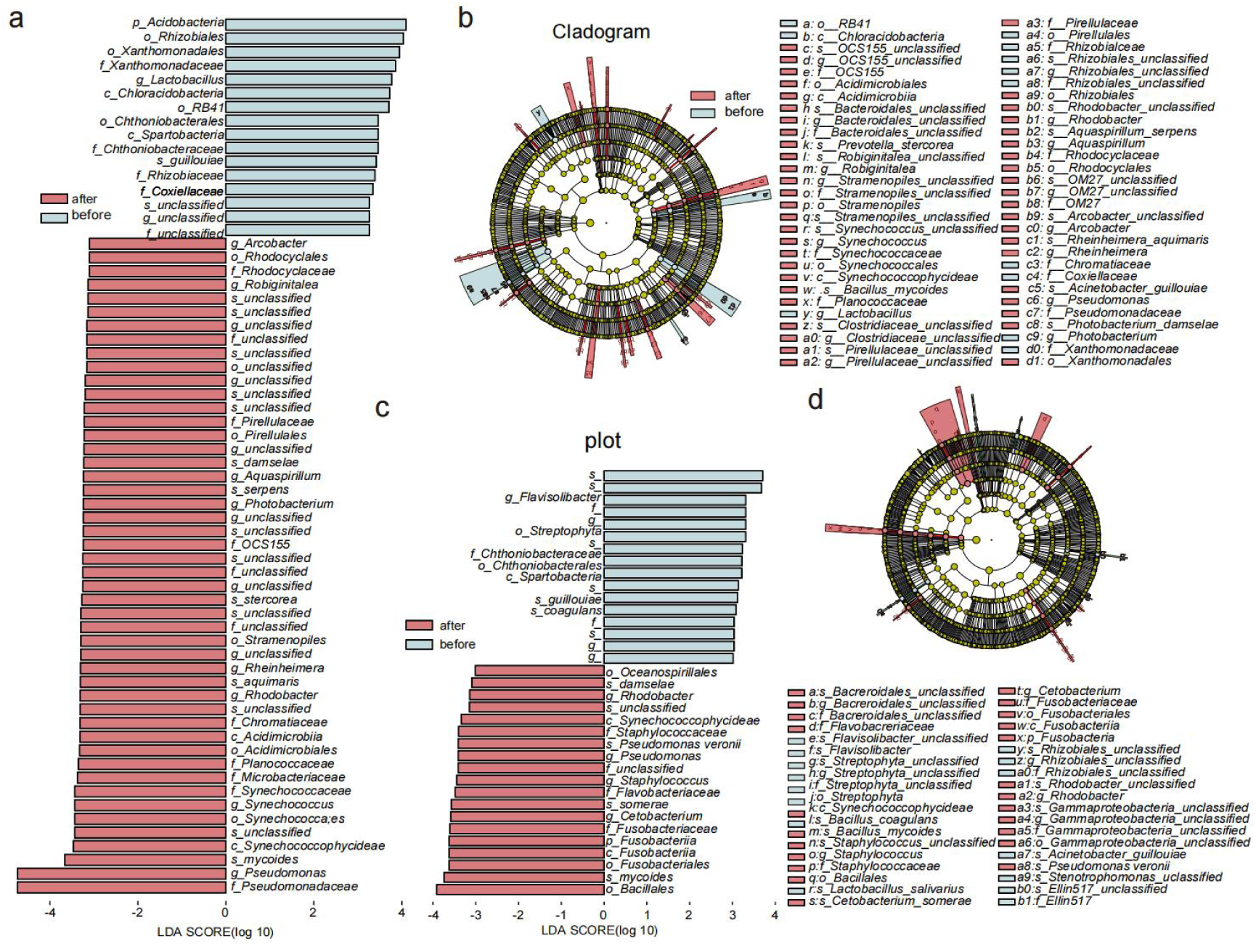
The diversity analysis of tumor-resident microbiota between pre- and post-drug resistance in lung cancer. **a**, Cladogram representation of data in pilot cohort (n=7). **b**, LEfSe analysis identifies the relative taxa abundance between pre- and post-drug resistance in pilot cohort. Taxa with LDA scores >2 are shown. **c**, Cladogram representation of data in validation cohort (n=114). **d**, LEfSe analysis identifies the relative taxa abundance between pre- and post-drug resistance in validation cohort (n=114). Taxa with LDA scores >2 are shown.

**Extended Data Fig. 2.**
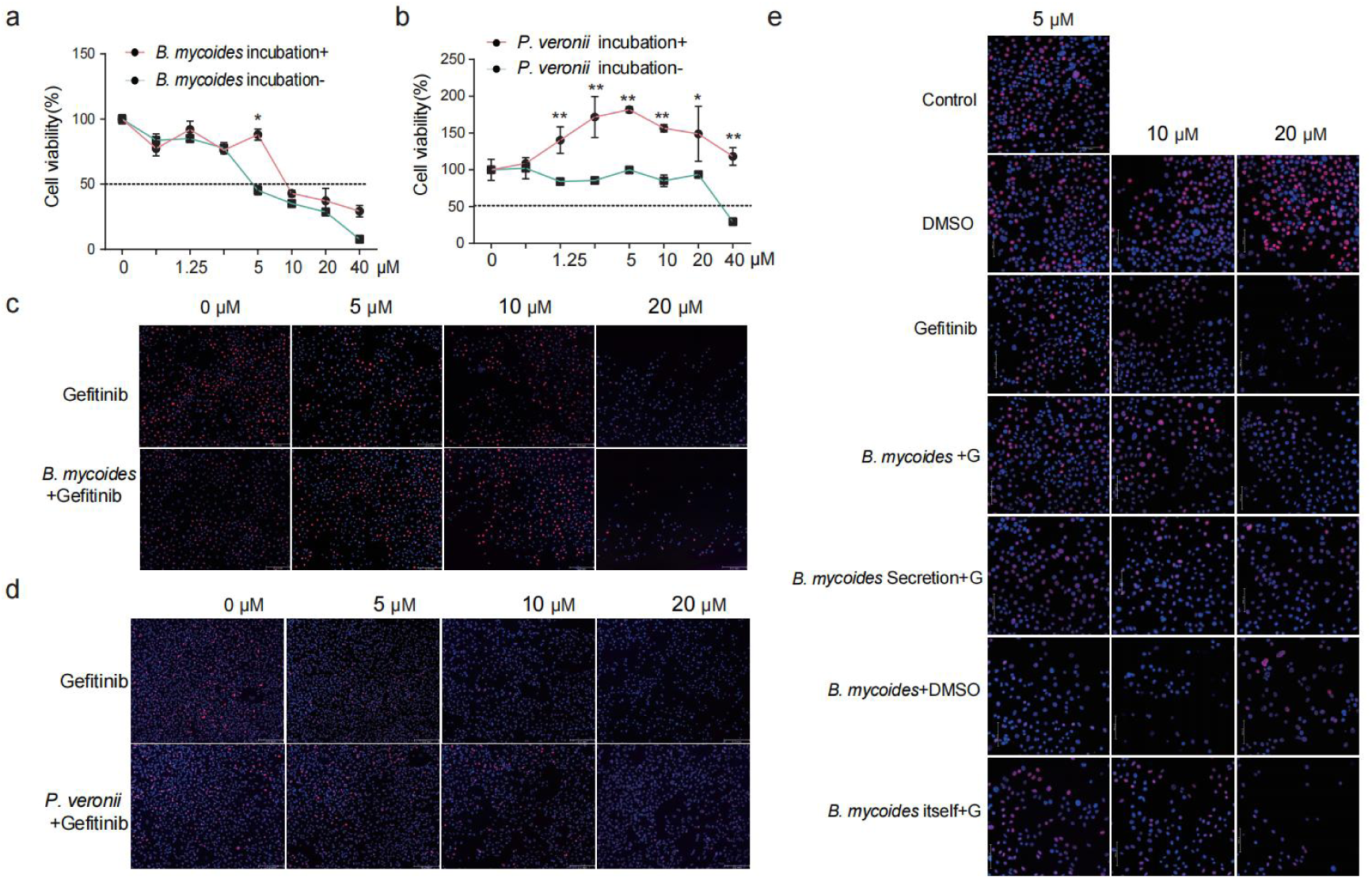
Gefitinib-tolerant cell lines demostrated *B. mycoides* and *P. veronii* could assist cells to tolerate Gefitinib in vitro. **a,b**, Relative proliferation rate of gefitinib-tolerant cell A549 administered with gefitinib preincubated with/without bacteria. *B. mycoides* (a) and *P. veronii* **(b)**. **c,d**, A549 cell proliferation was observed under a light microscope in response to gefitinib preincubated with/ without bacteria. Red flurescenece indicates EdU-positive cells stained with AZD555,blue flurescenece indicates cell nuclei stained with Hoechst 33342. *B. mycoides* (c) and *P. veronii* (d). **e**, A549 cell proliferation was observed under a light microscope in response to different treament conditions. *B. mycoides*, bacterial cells, or bacterial secretions preinbubated with gefitinib at 5μM,10μM,20μM.

**Extended Data Fig. 3.**
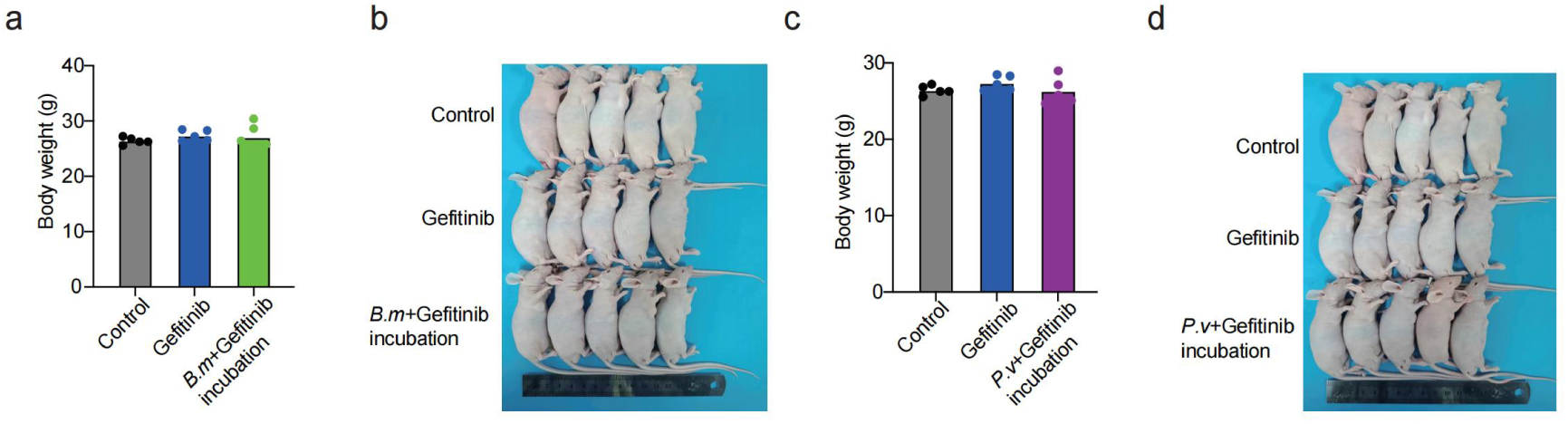
Gefitinib inhibit subcutaneous tumor growth but show no impact on mice weight. **a**, The body weight of mice under different treatment conditions. Gefitinib preincubatated with/without *B. mycoides.* **b**,The appearance of subcutaneous tumors in mice. The morphology and size of tumors under different treatment conditions with *B. mycoides.* **c**, The body weight of mice under different treatment conditions. Gefitinib preincubatated with/without *P. veronii.* **d**,The appearance of subcutaneous tumors in mice. The morphology and size of tumors under different treatment conditions with *P. veronii*.

**Extended Data Fig. 4.**
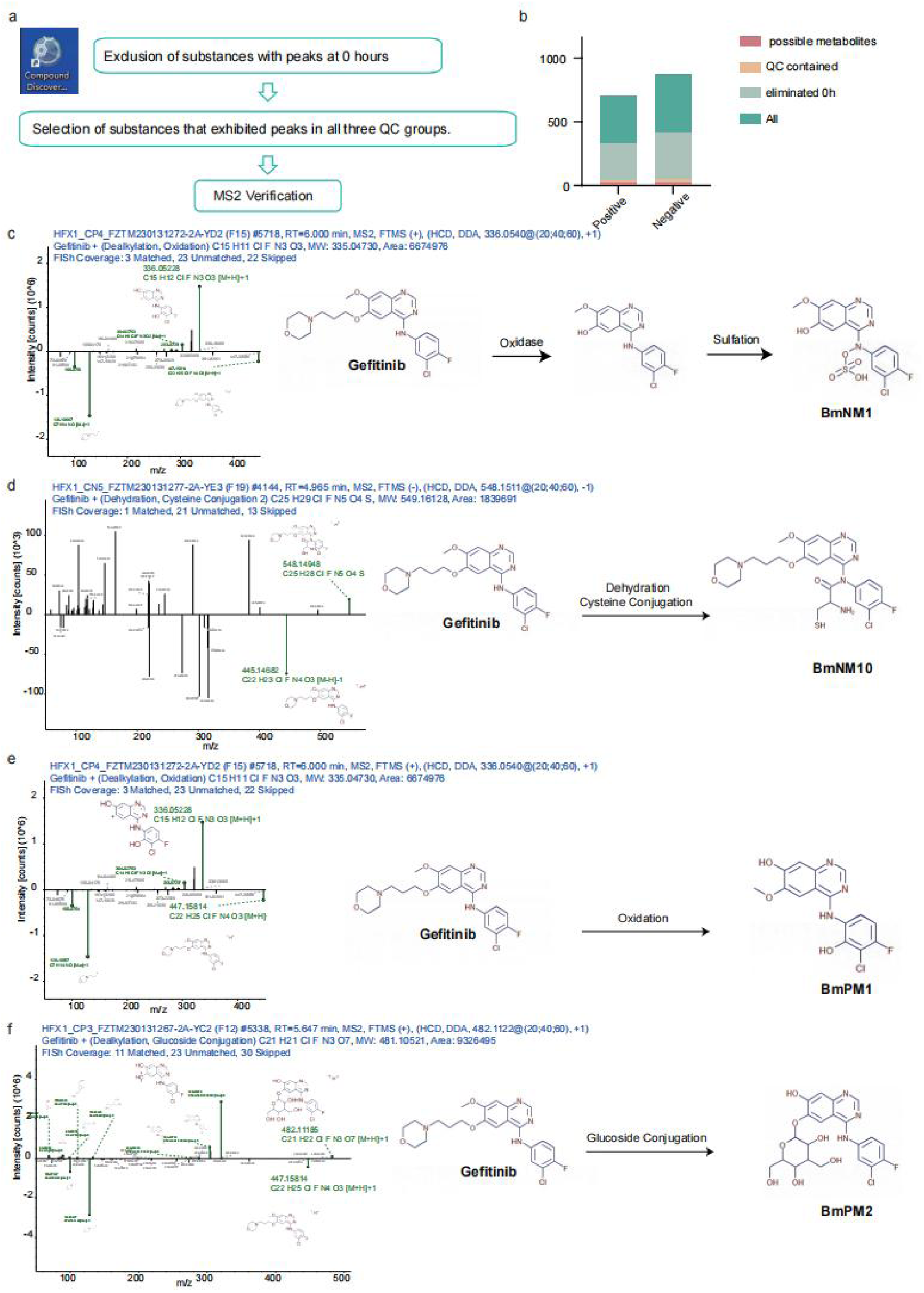
Elucidation of the structural characteristics of metabolites derived from the biotransformation of gefitinib by *B. mycoides*. **a**, Workflow diagram illustrating the metabolite identification process following database interrogation using Compound discoverer 3.1 data, with gefitinib as the parent compound. **b**, Distribution profile of metabolite abundance ratios across all mass spectrometry datasets. **c**, Extracted ion chromatogram of m/z 415.00411 and the proposed structure of BmNM1. **d**, Extracted ion chromatogram of m/z 549.16128 and the proposed structure of BmNM10. **e**, Extracted ion chromatogram of m/z 335.0473 and the proposed structure of BmPM1. **f**, Extracted ion chromatogram of m/z 481.10521 and the proposed structure of BmPM2.

**Extended Data Fig. 5.**
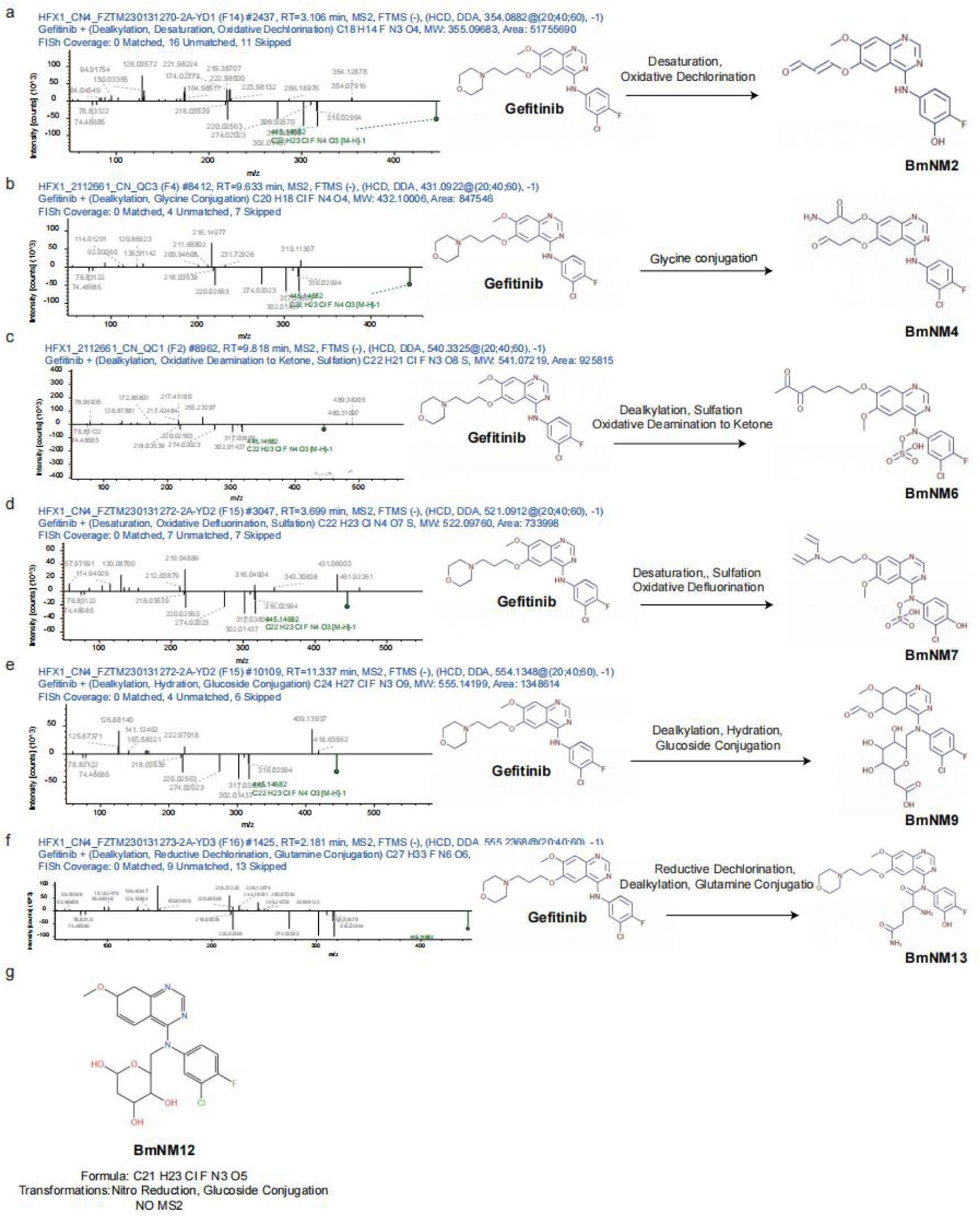
Structural characterization of biotransformation products derived from gefitinib by *B. mycoides* in negative ion mode mass spectrometry. **a**, Extracted ion chromatogram of m/z 355.09683 and the proposed structure of BmNM2. **b**, Extracted ion chromatogram of m/z 432.10006 and the proposed structure of BmNM4. **c**, Extracted ion chromatogram of m/z 541.07219 and the proposed structure of BmNM6. **d**, Extracted ion chromatogram of m/z 522.0976 and the proposed structure of BmNM7. **e**, Extracted ion chromatogram of m/z 555.14199 and the proposed structure of BmNM9. **f**, Extracted ion chromatogram of m/z 538.25398 and the proposed structure of BmNM13. **g**, The m/z 451.13103 and the proposed structure of BmNM5, No MS2 spectrum was detected for the compound.

**Extended Data Fig. 6.**
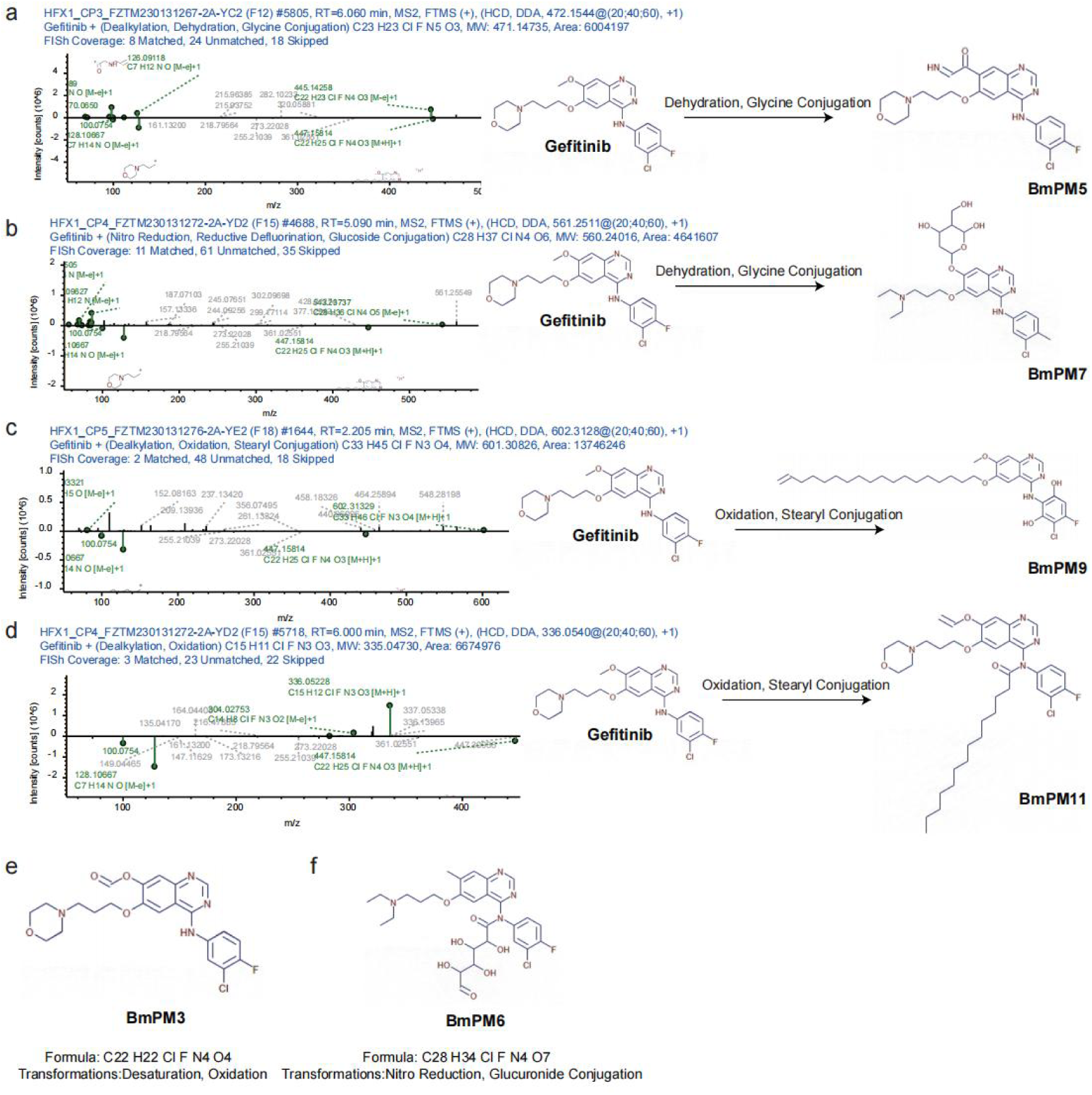
Structural characterization of biotransformation products derived from gefitinib by *B. mycoides* using positive ion mode mass spectrometry. **a**, Extracted ion chromatogram of m/z 471.14735 and the proposed structure of BmPM5. **b**, Extracted ion chromatogram of m/z 560.24016and the proposed structure of BmPM7. **c**, Extracted ion chromatogram of m/z 601.30826 and the proposed structure of BmPM9. **d**, Extracted ion chromatogram of m/z 699.34504 and the proposed structure of BmPM11. **e**, The m/z 460.13136 and the proposed structure of BmPM3. **f**, The proposed structure of BmPM6 corresponding to m/z 592.21001, for which also no MS2 spectrum was detected.

**Extended Data Fig. 7.**
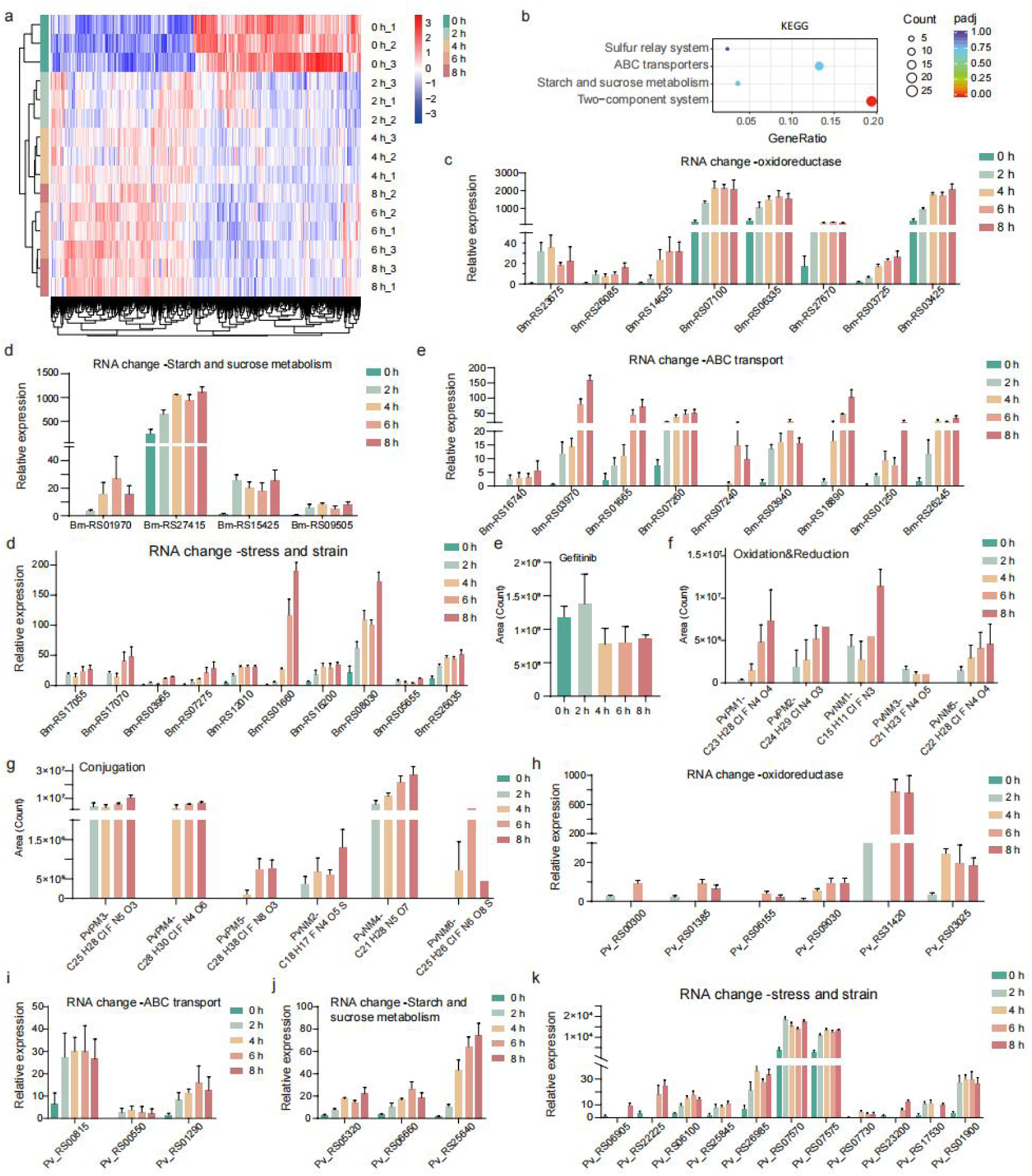
Temporal expression profiles of significantly upregulated genes in *B. mycoides* and temporal profiles of gefitinib metabolites and significantly upregulated genes responsive to gefitinib metabolism in *P. veronii*. **a**, Heatmap of transcriptional changes in all genes responsive to gefitinib metabolism in *B. mycoides.* **b**, KEGG pathway enrichment analysis of significantly upregulated genes responsive to gefitinib metabolism in *B. mycoides,* padj<0.01. **c**, Temporal expression profiles of significantly upregulated redox-related genes responsive to gefitinib metabolism in *B. mycoides,* padj<0.01. **d**, Temporal expression profiles of significantly upregulated starch and sucrose metabolism related genes responsive to gefitinib metabolism in *B. mycoides,* padj<0.01. **e**, Temporal expression profiles of significantly upregulated ABC transport system related genes responsive to gefitinib metabolism in *B. mycoides,* padj<0.01. **f**, Temporal expression profiles of significantly upregulated stress and strain related genes responsive to gefitinib metabolism in *B. mycoides,* padj<0.01. **g**, Time-dependent decrease in gefitinib concentration incubated with *P. veronii*. **h**, Time-course analysis of six potential gefitinib metabolites generated through redox metabolism by *P. veronii* using LC-MS. **i**, Time-dependent profiles of potential gefitinib conjugative metabolites generated by *P. veronii* using LC-MS. **j**, Temporal expression profiles of significantly upregulated redox-related genes responsive to gefitinib metabolism in *P. veronii,* padj<0.01. **k**, Temporal expression profiles of significantly upregulated ABC transport system related genes responsive to gefitinib metabolism in *P. veronii,* padj<0.01. **l**, Temporal expression profiles of significantly upregulated starch and sucrose metabolism related genes responsive to gefitinib metabolism in *P. veronii,* padj<0.01. **m**, Temporal expression profiles of significantly upregulated stress and strain related genes responsive to gefitinib metabolism in *P. veronii,* padj<0.01.

**Extended Data Fig. 8.**
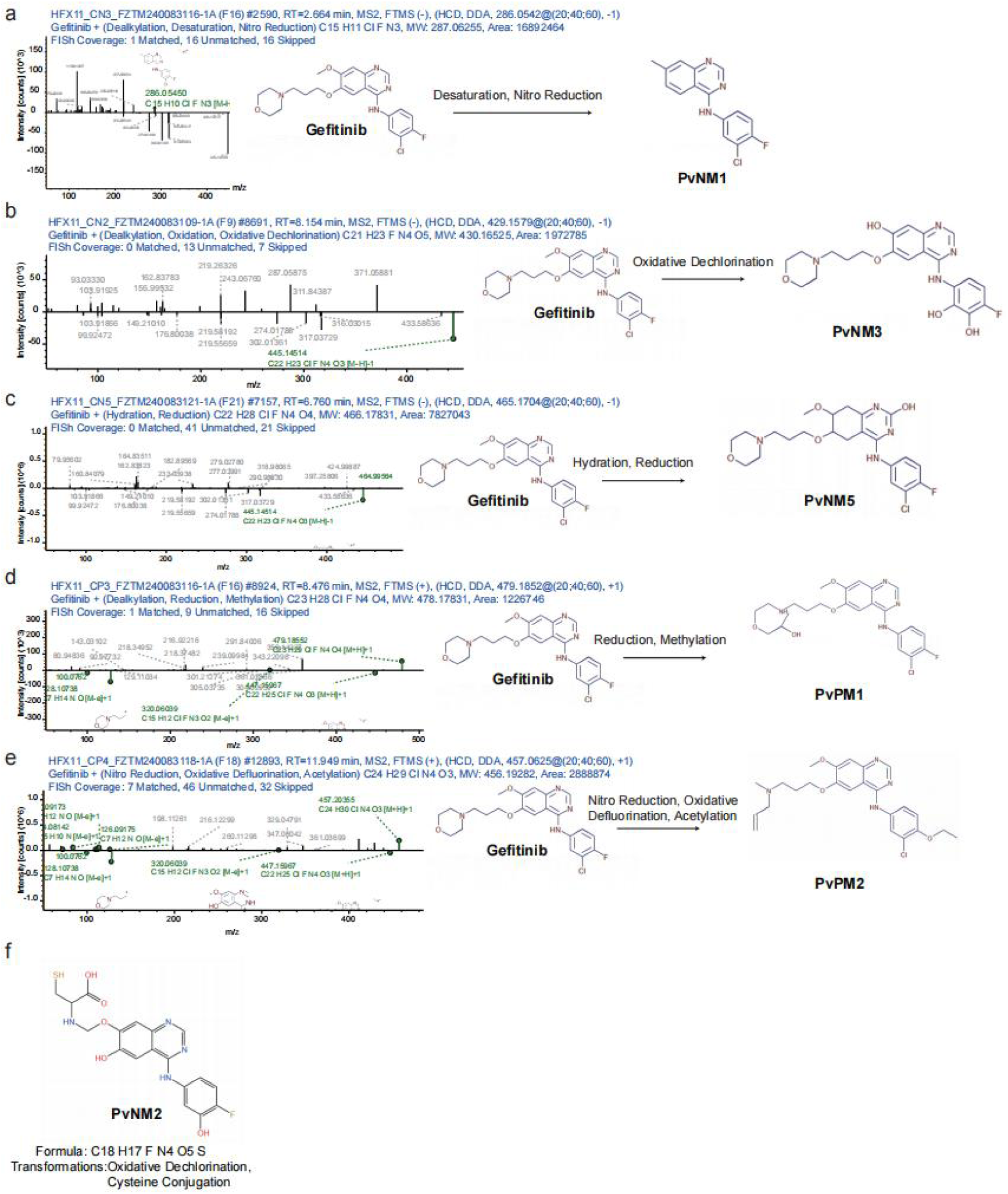
Structural characterization of biotransformation products derived from gefitinib by *P. veronii* using mass spectrometry. **a,** Extracted ion chromatogram of m/z 287.06255 and the proposed structure of PvNM1. **b**, Extracted ion chromatogram of m/z 430.16525 and the proposed structure of PvNM3. **c**, Extracted ion chromatogram of m/z 466.17831 and the proposed structure of PvNM5. **d**, Extracted ion chromatogram of m/z 478.17831 and the proposed structure of PvPM1. **e**, Extracted ion chromatogram of m/z 456.19282 and the proposed structure of PvPM2. **f**, The m/z 420.09037 and the proposed structure of PvNM2, No MS2 spectrum was detected for the compound.

**Extended Data Fig. 9.**
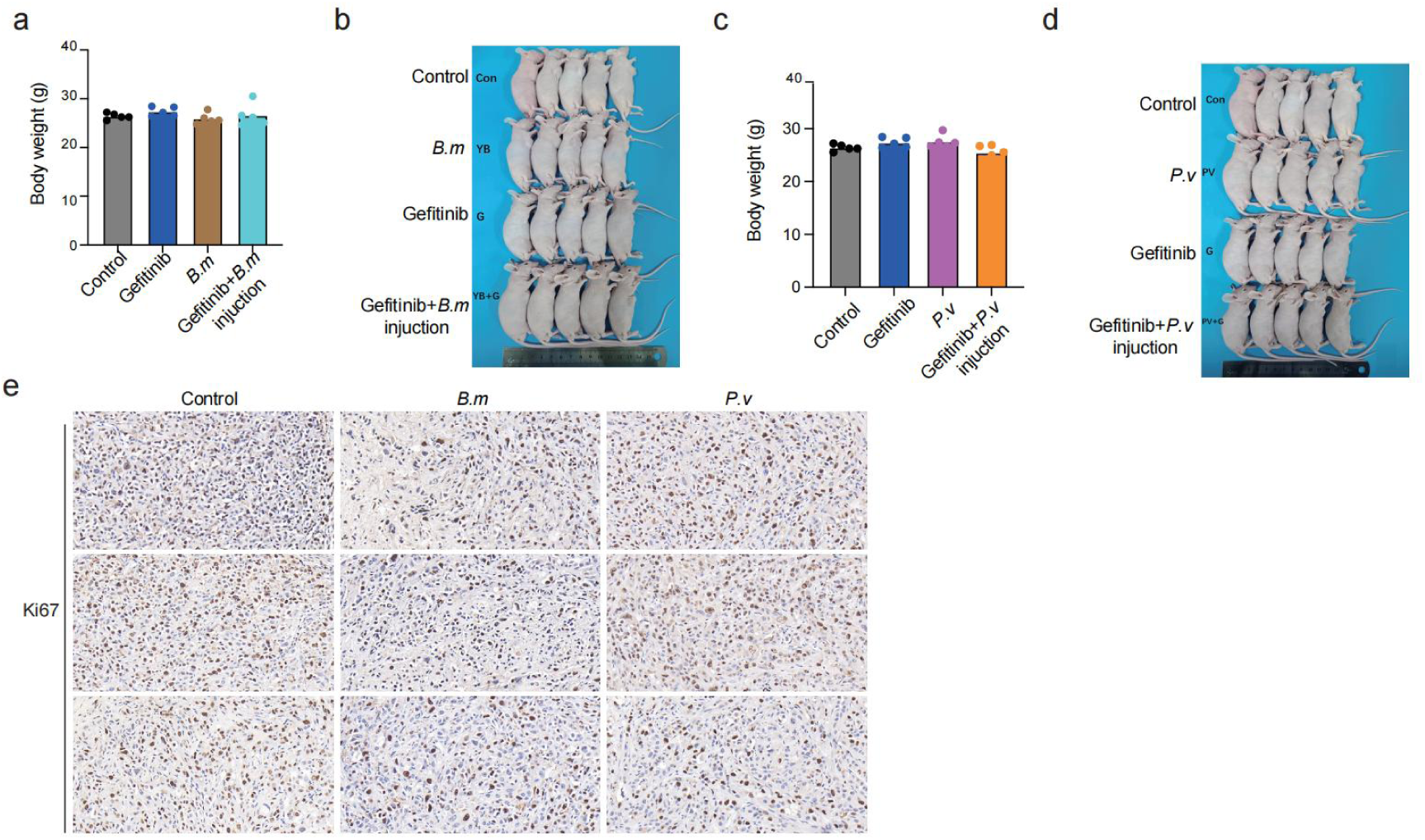
*B. mycoides* and *P. veronii* did not impact cell growth. **a**, The body weight of mice under different treatment conditions. *B. mycoides* was peritumoral injected while gefitinib was intraperitoneal injected. **b,** The appearance of subcutaneous tumors in mice. The morphology and size of tumors under different treatment conditions with *B. mycoides*. **c**, The body weight of mice under different treatment conditions. *P. veronii* was peritumoral injected while gefitinib was intraperitoneal injected. **d,** The appearance of subcutaneous tumors in mice. The morphology and size of tumors under different treatment conditions with *P. veronii*. **e**,Immunohistochemical stain of tumors stained with Rabbit anti-Ki67 when peritumoral injected with *B. mycoides* and *P. veronii*.

