## Supplementary figures and images for "Intratumoral Resident Microbes Directly Participate in Resistance to EGFR-TKI Targeted Drug Therapy through Metabolism in NSCLC"

### supplemental figure 1

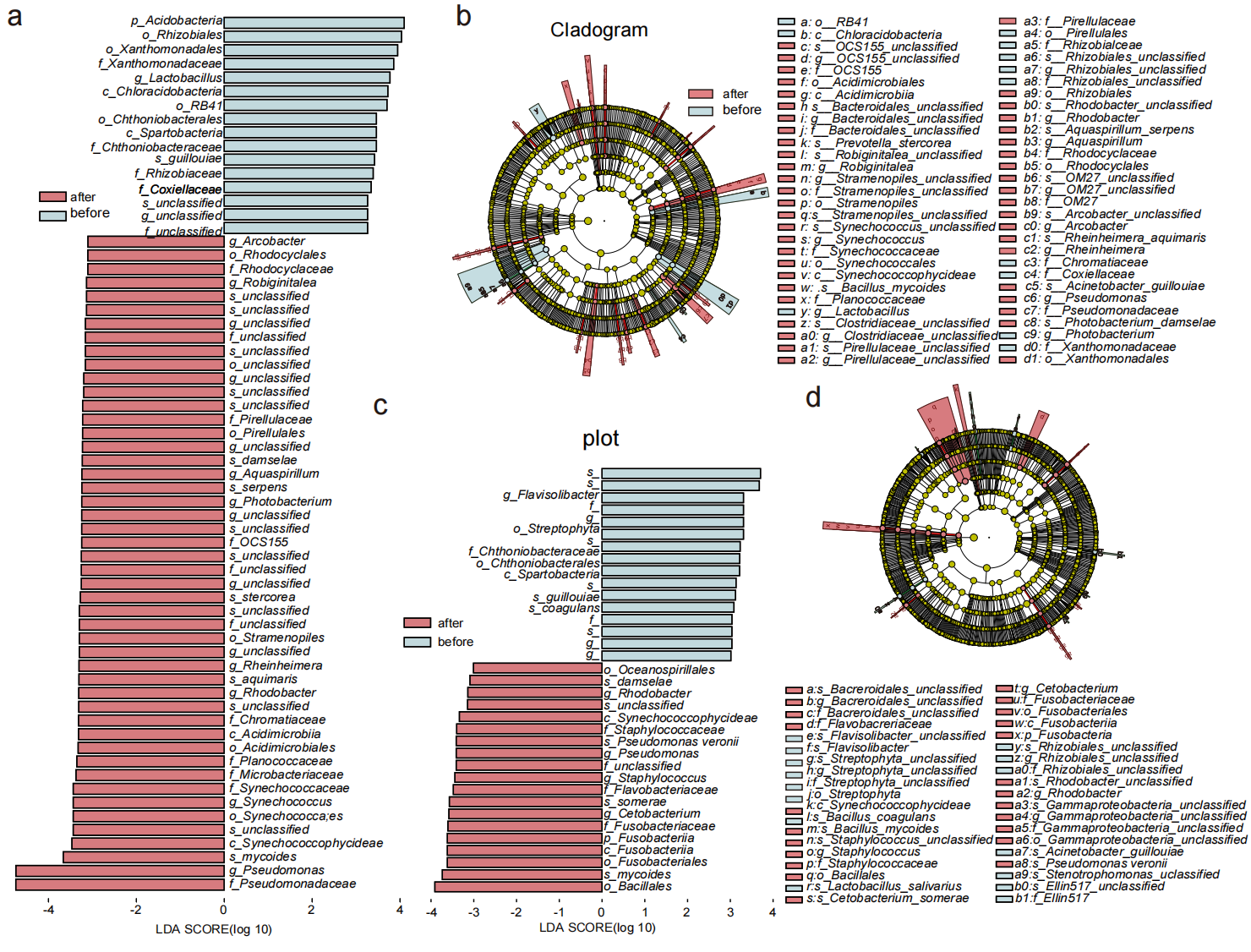

### supplemental figure 2

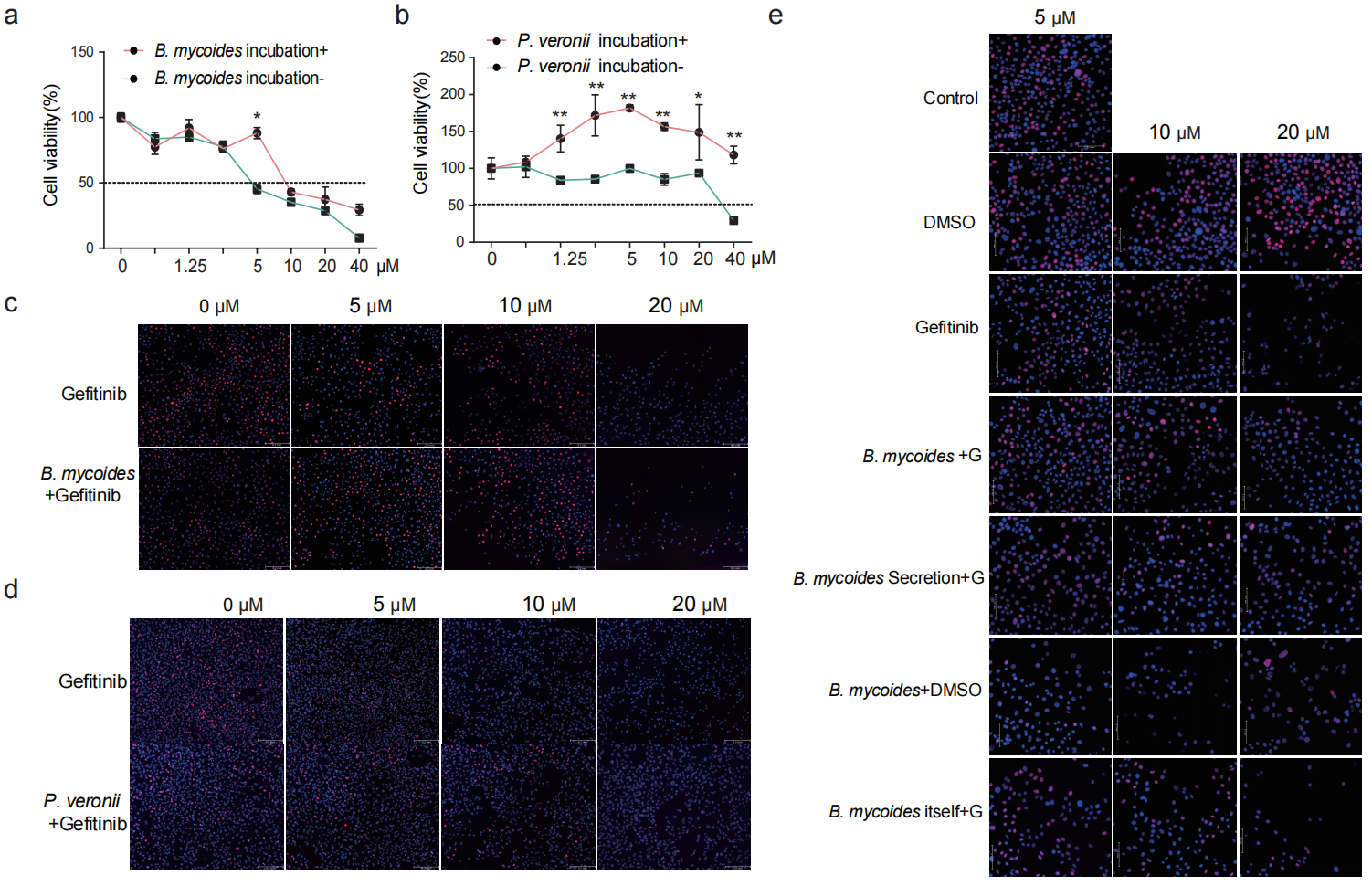

### supplemental figure 3

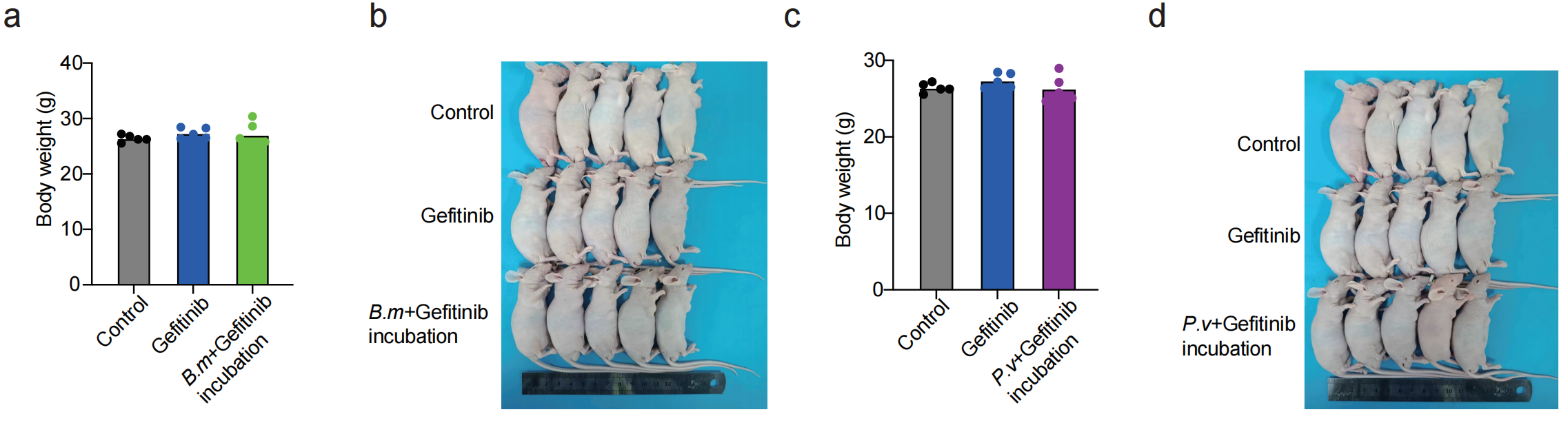

### supplemental figure 4

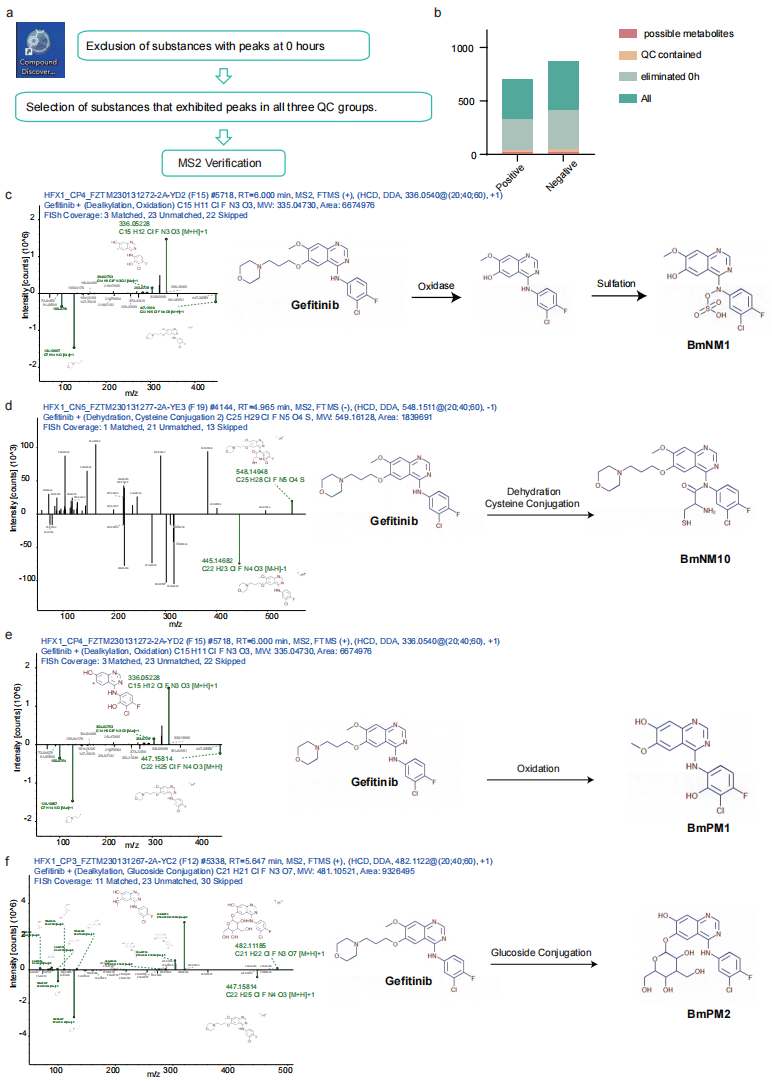

### supplemental figure 5

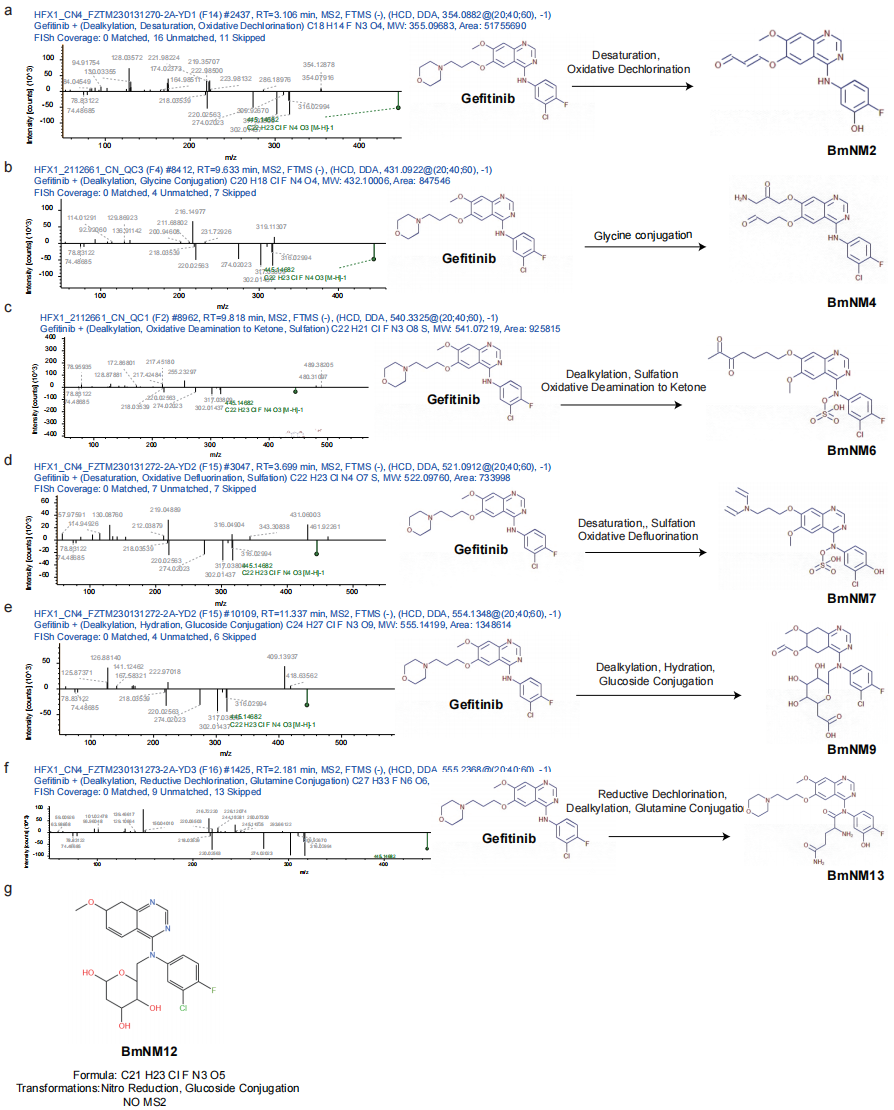

### supplemental figure 6

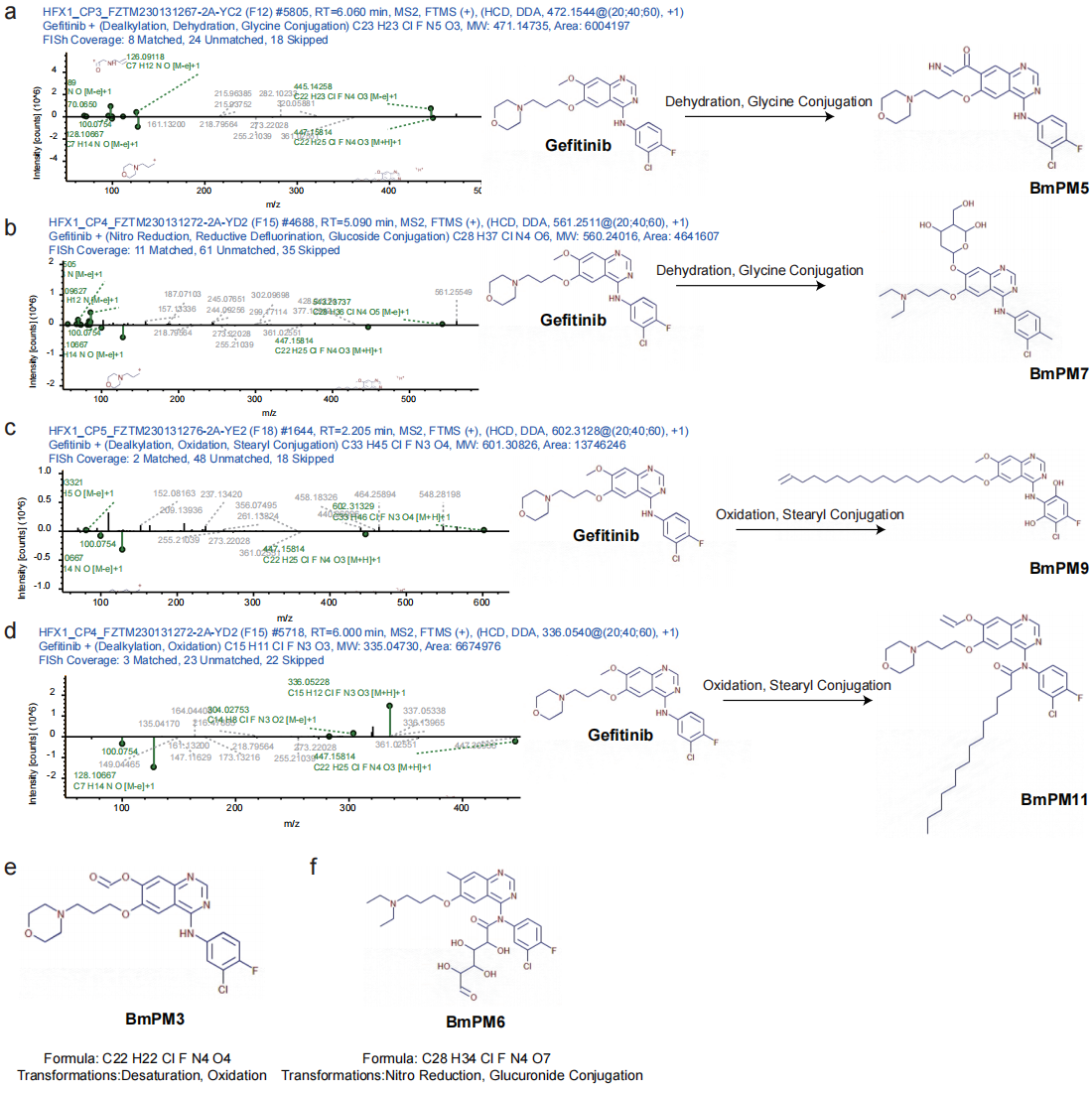

### supplemental figure 7

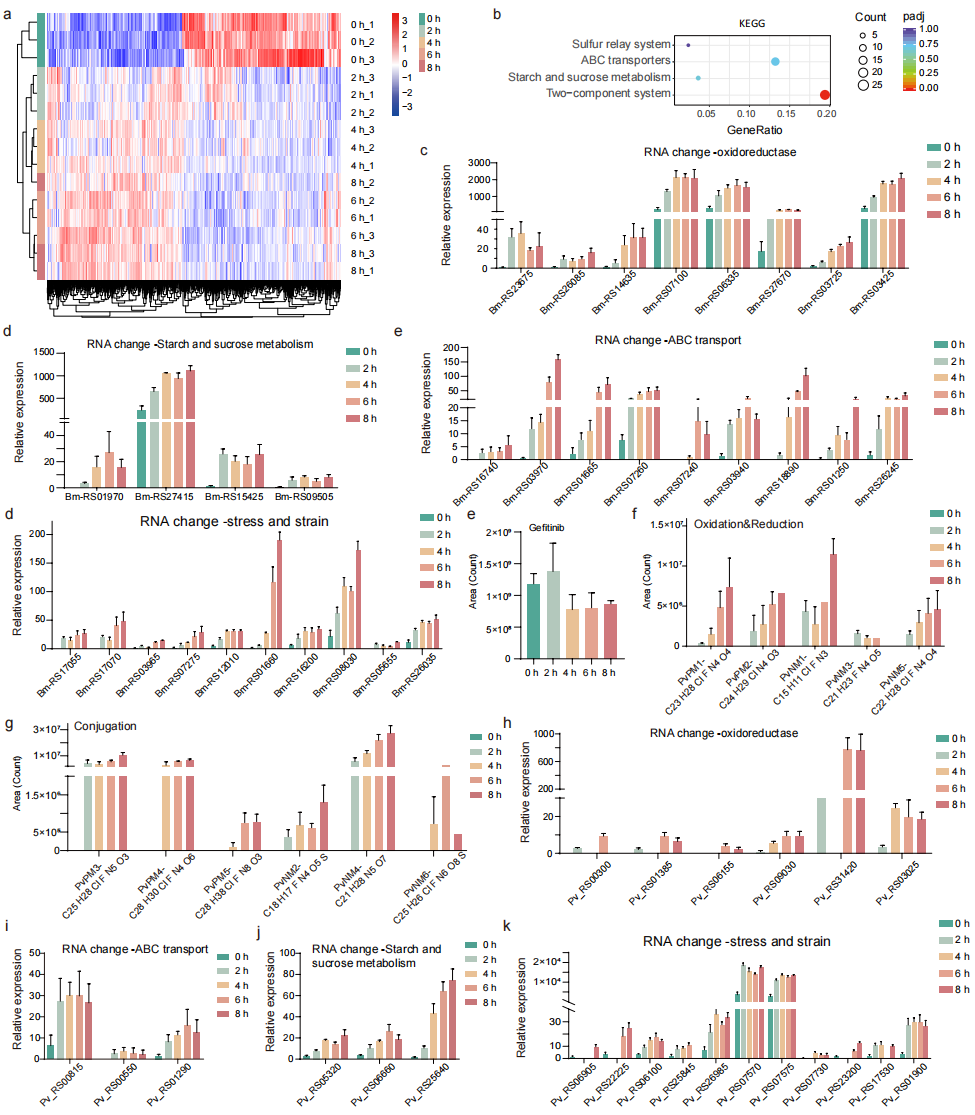

### supplemental figure 8

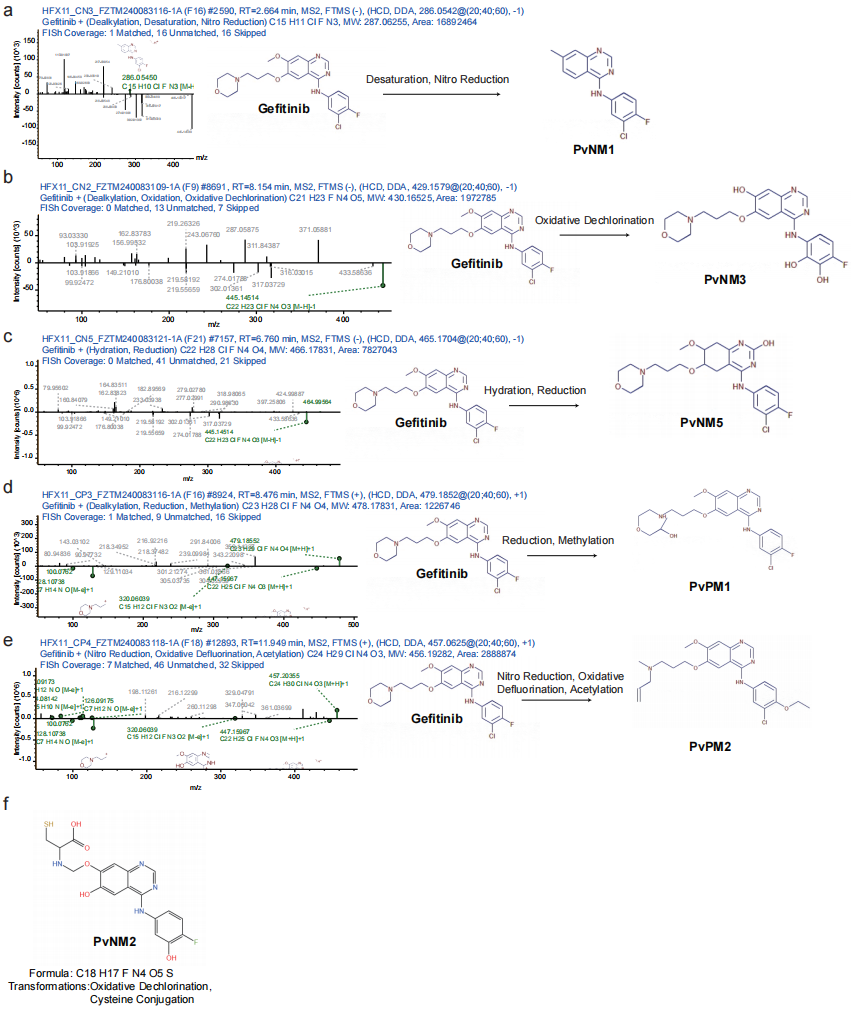

### supplemental figure 9

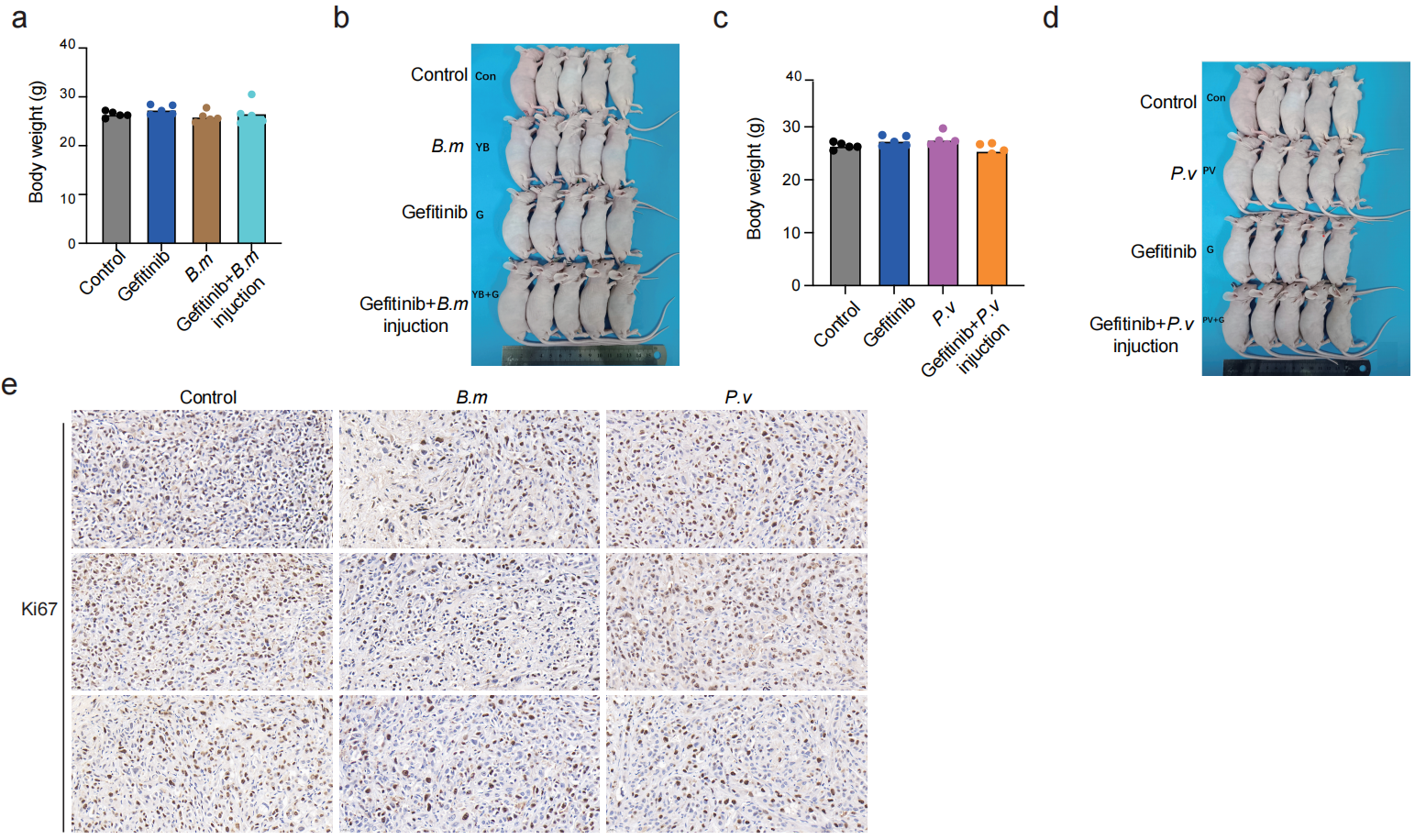
